# Complex host/symbiont integration of a multi-partner symbiotic system in the eusocial aphid *Ceratovacuna japonica*

**DOI:** 10.1101/2022.08.04.502717

**Authors:** Shunta Yorimoto, Mitsuru Hattori, Kondo Maki, Shuji Shigenobu

## Abstract

Some hemipteran insects rely on multiple endosymbionts for essential nutrients. However, the evolution of multi-partner symbiotic systems is not well-established. Here, we report a co-obligate symbiosis in the eusocial aphid, *Ceratovacuna japonica*. 16S rRNA amplicon sequencing unveiled co-infection with a novel *Arsenophonus* sp. symbiont and *Buchnera aphidicola*, a common obligate endosymbiont in aphids. Both symbionts were housed within distinct bacteriocytes and were maternally transmitted. The *Buchnera* and *Arsenophonus* symbionts had streamlined genomes of 432,286 bp and 853,149 bp, respectively, and exhibited metabolic complementarity in riboflavin and peptidoglycan synthesis pathways. These anatomical and genomic properties were similar to those of independently evolved multi-partner symbiotic systems, such as *Buchnera*–*Serratia* in Lachninae and *Periphyllus* aphids, representing remarkable parallelism. Furthermore, symbiont populations and bacteriome morphology differed between reproductive and soldier castes. Our study provides the first example of co-obligate symbiosis in Hormaphidinae and gives insight into the evolutionary genetics of this complex system.

## Introduction

Endosymbionts, microorganisms that live inside the body or cells of host organisms, are widespread in eukaryotes, serving as important resources for evolutionary innovation in hosts (Moran and Baumann, 2000; Nowack and Melkonian, 2010; Pawlowska et al., 2018; Rosenberg et al., 2010). Endosymbionts could confer beneficial functions to hosts, such as provisioning essential nutrients that are not synthesized by hosts (Akman Gündüz and Douglas, 2012; Douglas et al., 2001; Shigenobu et al., 2000; The International Aphid Genomics Consortium, 2010; Wilson et al., 2010) as well as providing resistance to heat stress (Chen et al., 2000; Russell and Moran, 2006) and protection against natural enemies via toxin production (Degnan et al., 2009; Nakabachi et al., 2013; Oliver et al., 2003). The acquisition of novel traits by endosymbionts contributes to niche divergence and lineage diversification of hosts (Janson et al., 2008; Sudakaran et al., 2017).

Aphids are good models for studying endosymbiosis. Almost all aphid species harbor the obligate mutualistic symbiont *Buchnera aphidicola* (Gammaproteobacteria) in specialized host cells known as bacteriocytes, and these symbionts are vertically transmitted to offspring (Braendle et al., 2003; Miura et al., 2003). *Buchnera* provides essential amino acids and riboflavin (vitamin B_2_), which aphids cannot synthesize and are deficient in plant phloem sap, a component of the aphid diet (Akman Gündüz and Douglas, 2012; Douglas et al., 2001; Nakabachi and Ishikawa, 1999; Shigenobu and Wilson, 2011). In addition to the primary symbiont *Buchnera,* aphids are often associated with facultative symbionts, which are not essential but can contribute to host fitness and phenotypes (Oliver et al., 2010). *Buchnera* genomes have experienced significant reductions down to ∼0.6 Mb with ∼600 protein-coding genes. Despite the drastic genome reduction, *Buchnera* genomes retain genes involved in the biosynthesis of amino acids and vitamins essential for the host aphids (Chong et al., 2019; Van Ham et al., 2003; Shigenobu et al., 2000). Facultative symbionts exhibit moderate genome reductions. For example, *Serratia symbiotica*, *Hamiltonella defensa* and *Regiella insecticola* of pea aphids have ∼2.0 Mb genomes encoding ∼2,000 genes (Burke and Moran, 2011; Degnan et al., 2009, 2010).

Recent genomic studies of aphid symbionts have revealed the dynamic evolution of coexisting bacterial partners that function collectively with the ancient and stable symbiont *Buchnera* (Shigenobu and Yorimoto, 2022). Novel symbionts, such as *Sphingopyxis*, *Pectobacterium*, *Sodalis*-related, and *Erwinia*-related bacteria, have been found by 16S ribosomal RNA amplicon sequencing in a wide variety of aphid lineages (McLean et al., 2019; Xu et al., 2020, 2021). In addition, co-obligate symbioses, where multiple species of symbionts are essential for host survival, have been found in several aphid lineages. Co-obligate symbiosis involving *Buchnera* and *Serratia* evolved repeatedly in multiple aphid lineages, including the subfamily Lachninae, genus *Periphyllus*, *Microlophium carnosum*, and *Aphis urticata* (Manzano-Marín et al., 2017; Monnin et al., 2020). Co-obligate *Buchnera* of Lachninae and *Periphyllus* (Chaitophorinae) aphids have ∼0.4 Mb genomes encoding ∼400 genes and have lost genes involved in the biosynthesis of some essential amino acids and vitamins that the co-obligate *Serratia* is presumed to compensate for (Lamelas et al., 2011a; Manzano-Marín et al., 2016; Monnin et al., 2020; Pérez-Brocal et al., 2006).

The tribe Cerataphidini (Hemiptera: Aphididae: Hormaphidinae) is an aphid clade that possesses a number of biologically interesting characteristics, including a complex life history, gall formation, and eusociality. They are able to alternate between primary host plants, where sexual reproduction occurs, and secondary host plants, where parthenogenesis occurs, seasonally (Aoki and Kurosu, 2010; Fukatsu et al., 1994). The fundatrix of Cerataphidini aphids hatches from an overwintering fertilized egg and induces morphologically diverse galls on the *Styrax* trees, their primary host plant (Aoki and Kurosu, 2010; Fukatsu et al., 1994). Cerataphidini aphids are eusocial; they produce sterile individuals known as a soldier caste to protect the colony from the natural enemies (Stern and Foster, 1996). Fukatsu et al. (1994) reported that several species of Cerataphidini have secondary symbionts in addition to *Buchnera* (Fukatsu et al., 1994), although their identities and roles remain elusive. In several Cerataphidini aphids, the primary symbiont *Buchnera* has been replaced with extracellular yeast-like symbionts (Fukatsu et al., 1994).

In this study, we investigated bacterial symbionts of *Ceratovacuna japonica* in the Cerataphidini tribe (Figure 1A), taking an advantage of a recently established laboratory rearing system for this species (Hattori et al., 2013). We uncovered two endosymbionts of *Ce. japonica* by deep sequencing of 16S rRNA amplicons, *Buchnera* and an *Arsenophonus*-related bacterium, and these were consistently detected together. We sequenced the whole genomes of the two symbionts to assess the roles of both symbionts. We conducted microscopic observations to investigate their localization and dynamics in the host. We also compared symbiont populations between social castes. Our results suggest that the newly detected *Arsenophonus*-related symbiont is a novel species that has established obligate relationships with the host *Ce. japonica* through collaboration with *Buchnera*.

**Figure 1.**
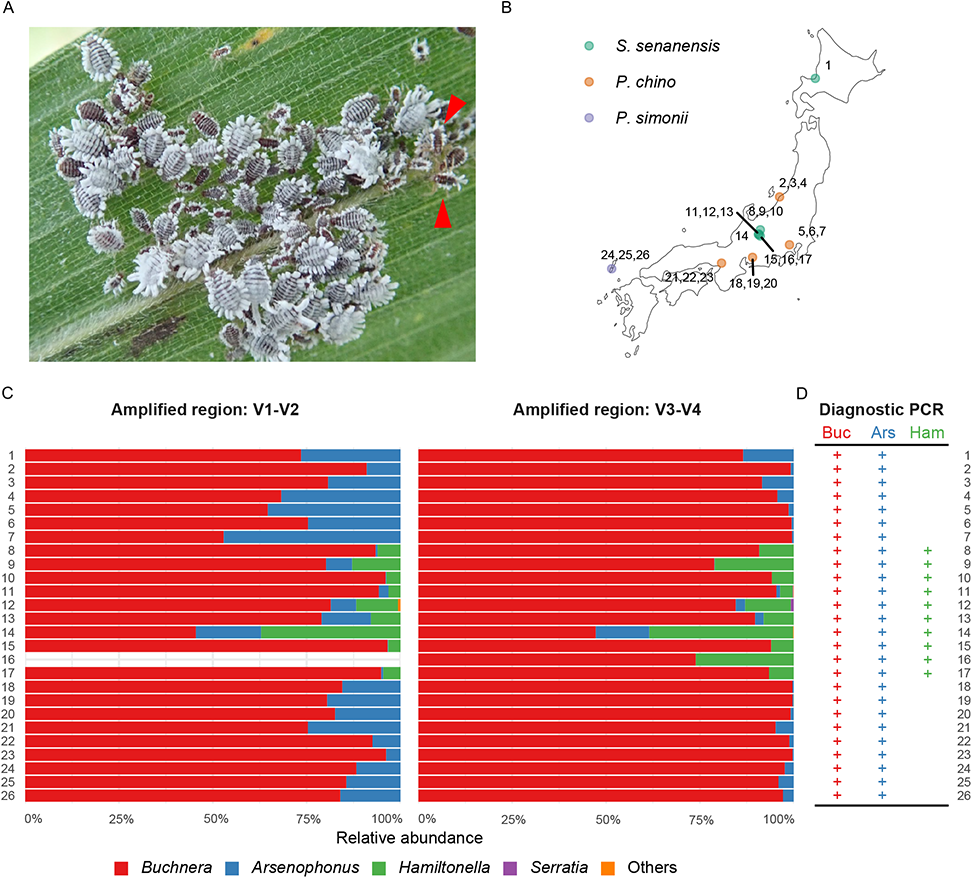
Symbiont composition determined by deep sequencing of 16S rRNA amplicons of natural populations of *Ceratovacuna japonica*. (A) A colony of *Ce. Japonica* on the bamboo grass *P. chino*. Red arrowheads indicate soldiers. (B) Locations and host plant species of collected *Ce. Japonica*. (C) Relative abundance of bacterial symbionts in *Ce. Japonica*. The V1–V2 region amplified from sample 16 was excluded owing to sequencing failure. “Others” includes unclassified sequences or those with low abundance. (D) Summary of diagnostic PCR detection of *Buchnera*, *Arsenophonus*, and *Hamiltonella* in *Ce. Japonica*. See also Figure S1 for agarose gel electrophoresis data. Each bacterial symbiont was detected using specific primers targeting a single-copy gene, *DnaK*. Each sample ID corresponds to those in Figure 1B, 1C, and S1. Detected bacterial symbionts are shown by the “+” character.

## Results

### Co-infection of *Buchnera* and *Arsenophonus* in natural populations of *Ceratovacuna japonica*

To assess the symbiont communities in *Ce. japonica*, we collected 26 colonies at nine geographically distinct populations from three different host plant species across Japan for high-throughput 16S rRNA amplicon sequencing (Figure 1B and 1C, Table S1 & S2). The sequencing of 16S rRNA hypervariable regions, V1–V2 and V3–V4, yielded 332,656 and 432,594 reads after quality filtering. V1–V2 and V3–V4 reads were classified into 13 and 25 OTUs (operational taxonomic units), respectively. The prevalent bacterial genera were identified as *Buchnera* (mean relative abundance ± SE of V1–V2: 81.48 ± 2.63%, V3–V4: 92.15 ± 2.22%), *Arsenophonus* (V1–V2: 14.89 ± 2.41%, V3– V4: 2.55 ± 0.76%), and *Hamiltonella* (V1–V2: 3.60 ± 1.58%, V3–V4: 5.25 ± 1.88%) (Figure 1C, Table S1 and S2). *Buchnera*, the nearly ubiquitous endosymbiont of aphids, was detected in all populations at an extremely high relative abundance (86.92 ± 1.86%) (Figure 1C). In addition, *Arsenophonus* was detected in all populations of *Ce. japonica*, regardless of host plant and geographical location (Figure 1B and 1C, Table S1 and S2). *Hamiltonella* was detected in specimens from 10 populations in the Nagano area (#8-17 in Figure 1) feeding on *Sasa senanensis*; however, it was not detected in the Hokkaido population (#1 in Figure 1) on the same host plant (Figure 1B and 1C, Table S1 and S2). *Serratia* was detected in only a single colony at a low level (Figure 1B and 1C, Table S2). We then performed PCR analyses to confirm the presence or absence of *Buchnera*, *Arsenophonus*, and *Hamiltonella* in the *Ce. japonica* populations with specific primers targeting a single-copy gene, *dnaK*, for each bacterium. Consistent with the result of 16S rRNA amplicon sequencing, PCR analyses showed that both *Buchnera* and *Arsenophonus* were detected in all populations, while *Hamiltonella* was detected only in populations in the central Japan area (Figure 1D and S1A). Next, we investigated the infection status at the individual level. Diagnostic PCR analyses of each of the 12 individuals in four geographically distinct populations detected both *Buchnera* and *Arsenophonus* in all cases (Figure S1B). Taken together, *Ce. japonica* was always co-infected with *Buchnera* and *Arsenophonus*, forming a multi-partner symbiosis. Additionally, *Ce. japonica* could be infected with *Hamiltonella* as a facultative symbiont.

We established an isofemale line of the aphid *Ce. japonica*, named NOSY1, from a single individual collected at the foot of Mt. Norikura (location #14 in Figure 1, S1 and Table S1–S2). It was cultured on bamboo shoots in the laboratory to undergo parthenogenetic reproduction for detailed molecular and microscopic analyses, as described below.

### Molecular phylogeny indicates *Arsenophonus* of *Ce. japonica* is related to obligate *Arsenophonus* symbionts of hematophagous insects

We performed molecular phylogenetic analyses based on the 16S rRNA sequences with our assembled data. *Buchnera* of *Ce. japonica* (hereafter referred to as *Buchnera* CJ for simplicity) was included within the *Buchnera* clade in Cerataphidini (Figure 2A). *Buchnera* CJ was closely related to a species in the same genus, *Ceratovacuna nekoashi*, and formed a sister group to *Ceratovacuna*, *Pseudoregma panicola* and *Pseudoregma bambucicola*. *Buchnera* of Nipponaphidini, Hormaphidini, and Cerataphidini species were monophyletic with clear separation between clades. These phylogenetic relationships of *Buchnera* mirrored those of their host aphid species (Chen et al., 2014; Ortiz-Rivas and Martínez-Torres, 2010).

**Figure 2.**
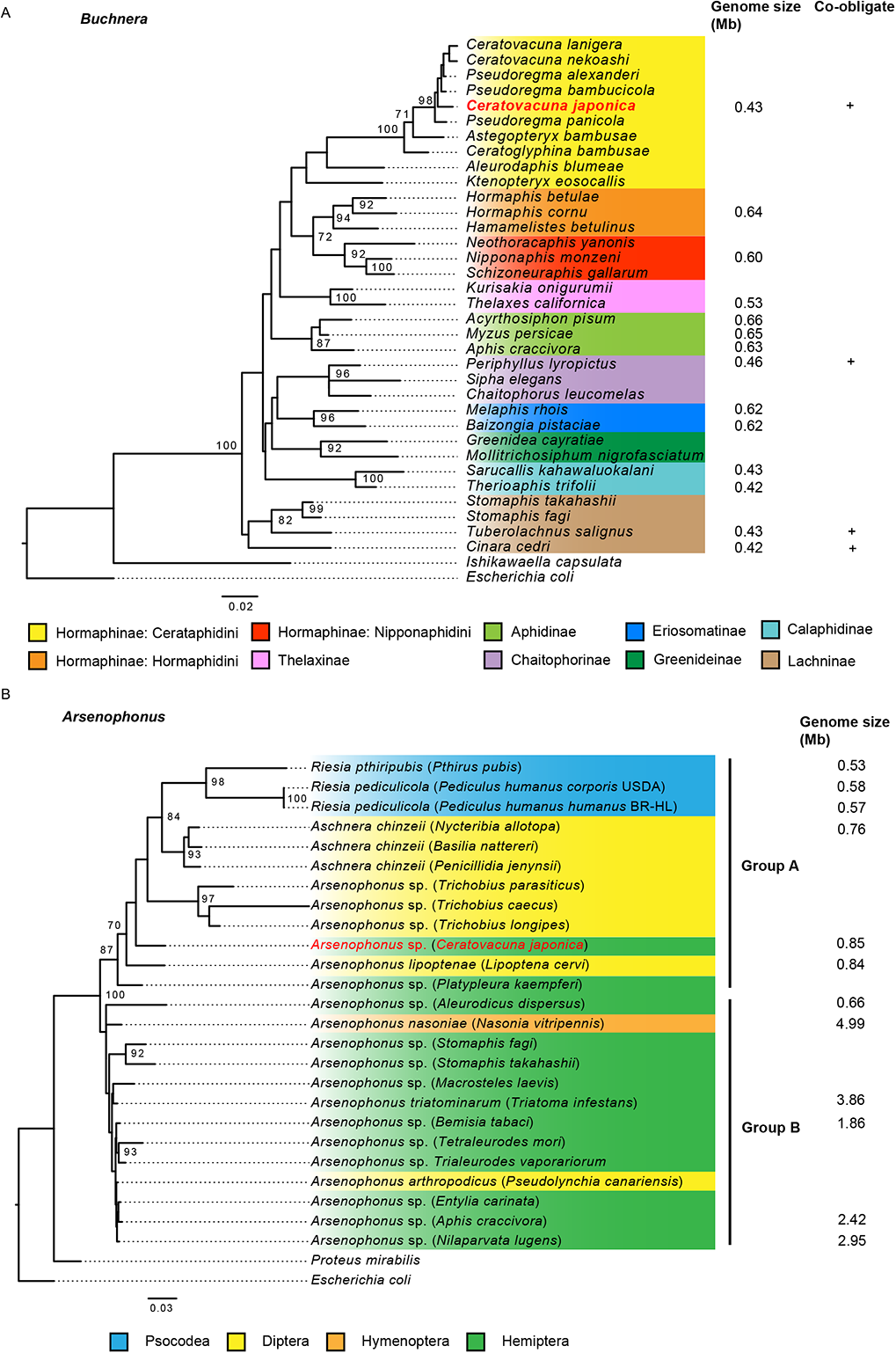
Phylogenetic analysis of *Buchnera* and *Arsenophonus* based on 16S ribosomal RNA sequences. (A) Maximum likelihood (ML) tree of *Buchnera aphidicola*. *Escherichia coli* and *Ishikawaella capsulata* were used as outgroups. The labels indicate the name of the host species. (B) ML tree of *Arsenophonus*. *E. coli* and *Proteus mirabilis* were used as outgroups. Target symbionts in this study are highlighted in red on each tree. Bootstrap values no less than 70% are indicated on each node. Scale bars represent 0.02 and 0.03 substitutions per site.

The 16S rRNA sequence of *Arsenophonus* of *Ce. japonica* (hereafter referred to as *Arsenophonus* CJ) showed the highest similarity (93.0%) to that of *Candidatus* Arsenophonus lipoptenae (Figure S3A), an obligate endosymbiont of the blood-sucking deer fly *Lipoptena cervi* with a role as a B vitamin provider (Nováková et al., 2016). In the phylogenetic analysis, *Arsenophonus* of *Ce. japonica* was related to endosymbionts of hematophagous Diptera (the superfamily Hippoboscoidea), such as *Lipoptena*, *Trichobius*, and lice species, named Group A. On the other hand, *Arsenophonus* CJ was not included in the clade of symbionts of Hemiptera (Group B in Figure 2B). The 16S rRNA sequence identity between *Arsenophonus* CJ and the closest relative, *Candidatus* Arsenophonus lipoptenae (i.e., 93.0%), was far lower than typical thresholds for assignment to the same species (e.g., ≤ 97%) (Stackebrandt and Goebel, 1994), indicating that the symbiont discovered in this study was a novel species in the genus *Arsenophonus*.

The 16S rRNA sequence of *Hamiltonella* of *Ce. japonica* (hereafter referred to as *Hamiltonella* CJ) was almost identical (99.8%) to that of *Candidatus* Hamiltonella defensa, a facultative symbiont of the pea aphid *Acyrthosiphon pisum* (Figure S3B).

### *Buchnera* and *Arsenophonus* are intracellular symbionts housed inside the distinct bacteriocytes

We inspected the internal morphology of *Ce. japonica* to uncover the localization of the three bacteria identified by 16S rRNA amplicon sequencing. Our histological observation identified a pair of symbiotic organs (bacteriomes) in the thorax of the adult parthenogenetic viviparous female (Figure 3AC). Each bacteriome exhibited an oval-shaped structure with a major axis of approximately 191 µm composed of several large presumably polyploid nuclei surrounded by the cytoplasm, which was filled with bacterial cells (Figure 3B), a typical characteristic of aphid bacteriocytes. Interestingly, HE staining and DAPI staining patterns allowed us to distinguish between two types of bacteriocytes. There were several uninucleate bacteriocytes on the surface of the bacteriome and a single double-nucleate bacteriocyte at the center of the bacteriome; the latter type of bacteriocytes showed less intense HE signals and more intence DAPI signals than those of the former type (Figure 3D and S3).

**Figure 3.**
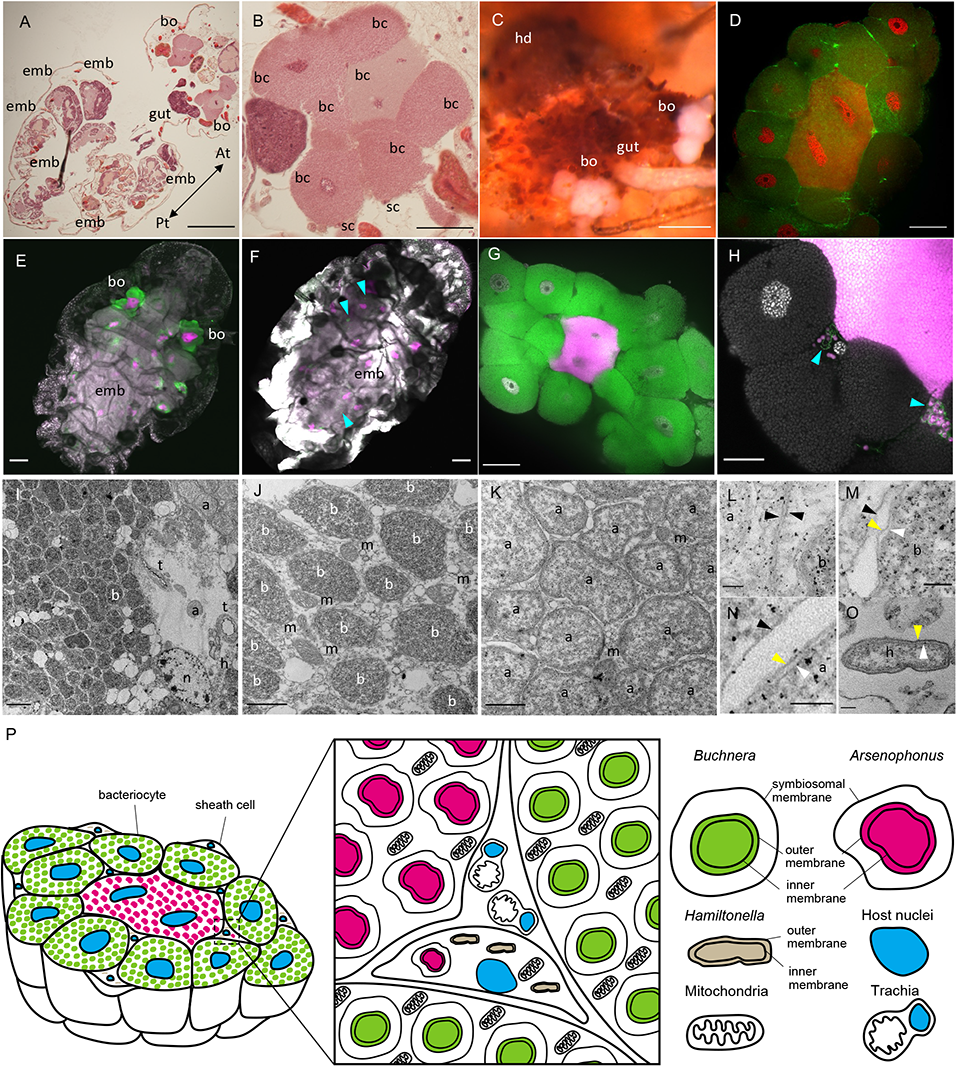
Localization and morphology of *Buchnera* and *Arsenophonus* symbionts. (A and B) Light microscopic images of tissue sections stained with hematoxylin and eosin. Lateral sections of an adult individual are shown. The whole body (A) and the magnified image of the bacteriome (B). (C) Light microscopic image of dissected bacteriomes. (D) Confocal image of the bacteriome structure. Nuclei (red) and F-actin (green) were stained by DAPI and phalloidin. (E–H) Confocal images of *Buchnera*, *Arsenophonus,* and *Hamiltonella* stained with fluorescent probes specific to each bacterium. Whole bodies of adult individuals (E and F) and dissected bacteriomes (G and H). In (E and G), gray (DAPI), green (Cy5), and magenta (Cy3) signals indicate nuclei, *Buchnera,* and *Arsenophonus*, respectively. In (F and H), gray (DAPI), green (Cy5), and magenta (Cy3) signals indicate nuclei, *Hamiltonella,* and *Arsenophonus*, respectively. Cyan arrowheads indicate *Hamiltonella* in (F and H). (I–O) Electron microscopy of a bacteriome dissected from an adult individual. (I) Low-magnification image of three types of symbionts in the bacteriome. (J and K) Cytoplasm of a bacteriocyte harboring *Buchnera* (J) and *Arsenophonus* (K). (L) Boundary of bacteriocytes harboring *Buchnera* and bacteriocytes harboring *Arsenophonus*. Black arrowheads indicate membranes of bacteriocytes. (M–O) High magnification images of membranes of *Buchnera* (M), *Arsenophonus* (N), and *Hamiltonella* (O). In (M–O), black, yellow, and white arrowheads indicate the symbiosomal membrane, outer membrane, and inner membrane, respectively. (P) Schematic diagram of the bacteriome morphology. Scale bars show 200 µm in (A and C), 100 µm in (E and F), 50 µm in (B, D, and G), 20 µm in (H), 2 µm in (I), 1 µm in (J and K), and 100 nm in (L–O). a, *Arsenophonus*; b, *Buchnera*; bc, bacteriocyte; bo, bacteriome; emb, embryo; h, *Hamiltonella*; m, mitochondria; n, host nucleus; sc, sheath cell; t, trachea.

We next conducted fluorescence *in situ* hybridization (FISH) using probes specifically targeting each bacterium, *Buchnera* CJ, *Arsenophonus* CJ, and *Hamiltonella* CJ. The FISH analysis revealed that *Buchnera* and *Arsenophonus* infected the same bacteriome but were localized in different bacteriocytes, consistent with the differential staining pattern of HE and DAPI. *Buchnera* was localized in the uninucleate bacteriocytes in the outer layer of the bacteriome, while *Arsenophonus* was localized in the single double-nucleate bacteriocyte at the center of the bacteriome (Figure 3E and 3G). Both *Buchnera* and *Arsenophonus* were observed exclusively in these specialized bacteriocytes and we had no evidence for the extracellular localization (e.g. hemolymph) of these endosymbionts (Figure 3E). In addition, ovarioles in the abdomen contained many embryos infected with both *Buchnera* and *Arsenophonus* (Figure 3E), suggesting that both symbionts are maternally and vertically transmitted to offspring. In contrast to the systematic localization of *Buchnera* and *Arsenophonus* in the bacteriocytes, sporadic *Hamiltonella* were found in the hemocoel and some were detected in the sheath cells, where they often coexisted with *Arsenophonus* (Figure 3F and 3H).

Electron microscopy also revealed three types of symbionts that differed in morphology and electron density (Figure 3I). *Buchnera* and *Arsenophonus* were round and resided within their specialized bacteriocytes, whereas *Hamiltonella* was rod-shaped and resided within sheath cells (Figure 3I-3K and 3O). The *Buchnera*-containing bacteriocytes harbored more mitochondria in the cytoplasm than the *Arsenophonus*-containing bacteriocytes (Figure 3J and 3K). *Buchnera* and *Arsenophonus* had a three-layered membrane structure (Figure 3M and 3N), indicating that both symbionts are packed inside the host-derived membranes called symbiosomal membranes, as reported in *Buchnera* of the pea aphid (Munson et al., 1991). In contrast, *Hamiltonella* had only a two-layered membrane, i.e., a bacterial inner membrane and outer membrane (Figure 3O).

### Streamlined small genomes of *Buchnera* CJ and *Arsenophonus* CJ exhibit complementary metabolic capacity

We conducted a shotgun sequencing of the hologenome of the isofemale line of *Ce. japonica* NOSY1. A metagenomic assembly approach (see STAR methods for details) yielded assemblies of the complete genomes of three bacterial symbionts, *Buchnera* CJ, *Arsenophonus* CJ, and *Hamiltonella* CJ. The *Buchnera* CJ genome consisted of one circular chromosome and two plasmids, pLeu and pTrp. (Figure 4A, D and Table 1). The *Buchnera* CJ chromosome had a length of 414,725 bp, G+C content of 20.0%, and coding density of 88.5%. The genome size of *Buchnera* CJ was significantly smaller than the typical size (approximately 600 kb) of *Buchnera* of many aphid species (Chong et al., 2019; Van Ham et al., 2003; Shigenobu and Yorimoto, 2022; Shigenobu et al., 2000; Tamas et al., 2002). It was similar to the sizes of genomes of *Buchnera* of aphids belonging to the genera *Cinara*, *Tuberolachnus*, and *Periphyllus* (Lamelas et al., 2011a; Manzano-Marín et al., 2016; Monnin et al., 2020; Pérez-Brocal et al., 2006), all of which coexist with another obligate symbiont, *Serratia symbiotica*. The chromosome of *Buchnera* CJ encoded 370 protein-coding genes (CDSs), 3 rRNAs (16S, 5S, and 23S), 30 tRNAs, and 6 pseudogenes. The plasmid pLeu consisted of at least two tandem repeats, the 7,001 bp units of six genes (Figure 4D). The plasmid pTrp consisted of at least two tandem repeats, the 10,560 bp units of one *trpE*, one pseudogenized *trpG*, and 10 *trpG* genes (Figure 4D).

**Figure 4.**
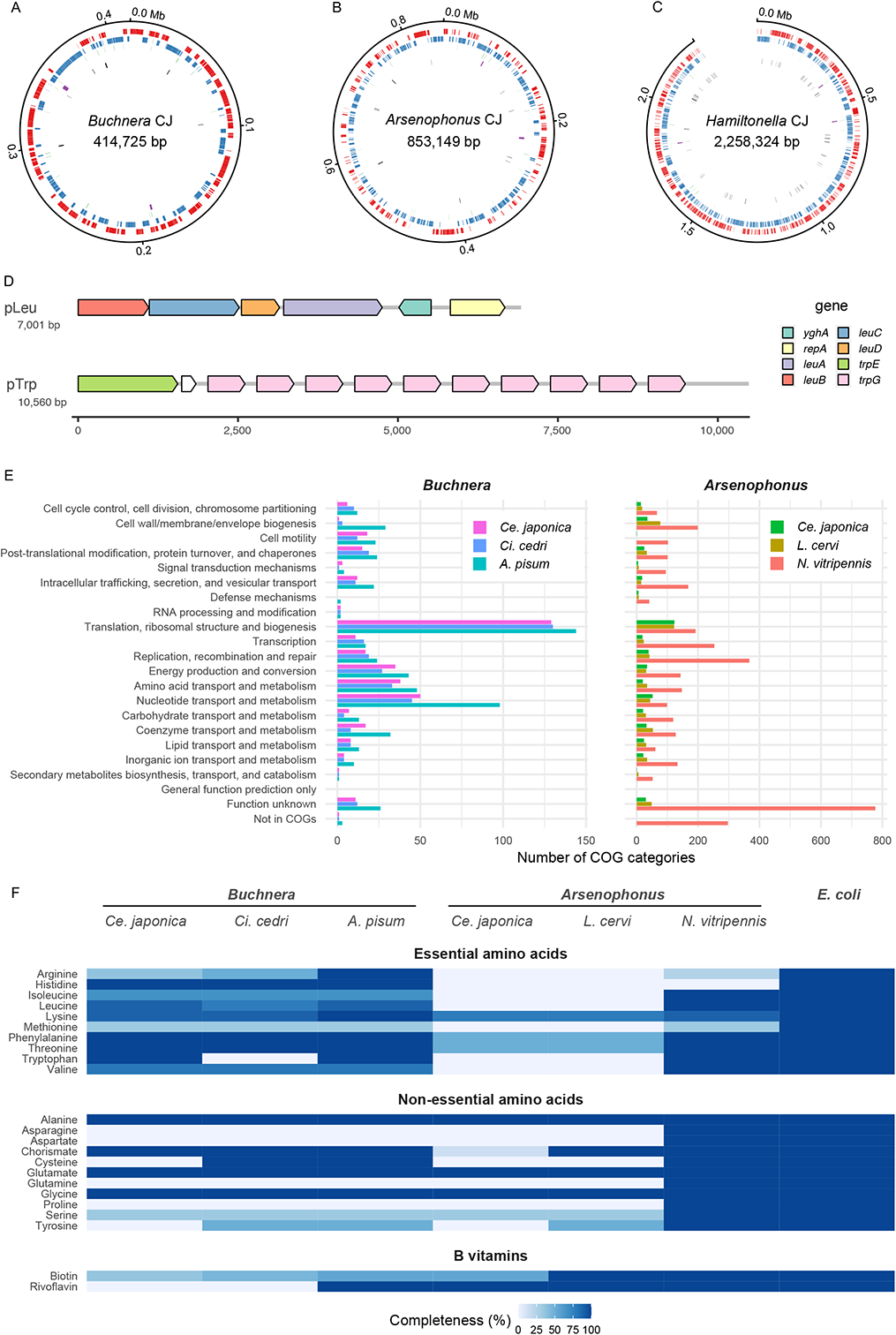
Genomic features of *Buchnera* CJ and *Arsenophonus* CJ. (A) Circular *Buchnera* CJ genome. (B) Circular *Arsenophonus* CJ genome. (C) Linear *Hamiltonella* CJ genome. Outer- to innermost rings correspond to (i) genome coordinates in kilobases; (ii) predicted protein-coding genes on the plus strand (red); (iii) predicted protein-coding genes on the minus strand (blue); (iv) transfer RNAs (green); (v) ribosomal RNAs (purple); (vi) pseudogenes (black) in (A–C). (D) Gene orders of plasmids of *Buchnera* CJ pLeu and pTrp. Arrows indicate the direction of transcription. White arrows indicate pseudogenes. (E) COG classification of protein-coding genes of *Buchnera* and *Arsenophonus*. (F) Comparison of gene repertoires responsible for nutrient synthesis by *Buchnera* and *Arsenophonus*. *E. coli* is shown as an example of a free-living bacterium. Gradient blue colors indicate the completeness of the minimal gene set for metabolic pathways.

**Table 1.**
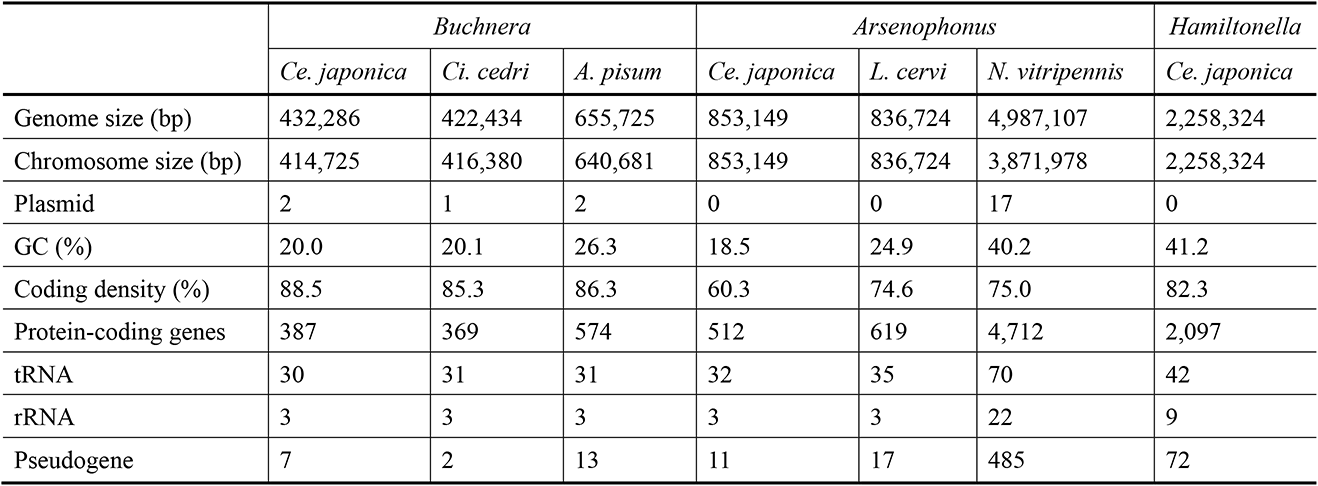
General features of sequenced symbiont genomes of *Ce. japonica* and the closely related species. *Buchnera aphidicola* of *Cinara cedri* (GCA_000090965.1); *B. aphidicola* of *Acyrthosiphon pisum* (GCA_000009605.1); *Candidatus* Arsenophonus lipoptenae of *Lipoptena cervi* (GCA_001534665.1); *Arsenophonus nasoniae* of *Nasonia vitripennis* (GCA_004768525.1).

|  | *Buchnera* |  |  | *Arsenophonus* |  |  | *Hamiltonella* |
| --- | --- | --- | --- | --- | --- | --- | --- |
|  | *Ce. japonica* | *Ci. cedri* | *A. pisum* | *Ce. japonica* | *L. cervi* | *N. vitripennis* | *Ce. japonica* |
| Genome size (bp) | 432,286 | 422,434 | 655,725 | 853,149 | 836,724 | 4,987,107 | 2,258,324 |
| Chromosome size (bp) | 414,725 | 416,380 | 640,681 | 853,149 | 836,724 | 3,871,978 | 2,258,324 |
| Plasmid | 2 | 1 | 2 | 0 | 0 | 17 | 0 |
| GC (%) | 20.0 | 20.1 | 26.3 | 18.5 | 24.9 | 40.2 | 41.2 |
| Coding density (%) | 88.5 | 85.3 | 86.3 | 60.3 | 74.6 | 75.0 | 82.3 |
| Protein-coding genes | 387 | 369 | 574 | 512 | 619 | 4,712 | 2,097 |
| tRNA | 30 | 31 | 31 | 32 | 35 | 70 | 42 |
| rRNA | 3 | 3 | 3 | 3 | 3 | 22 | 9 |
| Pseudogene | 7 | 2 | 13 | 11 | 17 | 485 | 72 |

The *Arsenophonus* CJ genome consisted of one circular chromosome (Figure 4B and Table 1). The *Arsenophonus* CJ chromosome had a length of 853,149 bp and G+C content of 18.5%. The genome of *Arsenophonus* CJ was considerably smaller than the genome of *Arsenophonus nasoniae* (5.0 Mb), a son-killer bacterium of *Nasonia vitripennis* (Darby et al., 2010), and similar in size to the genome of *Candidatus* Arsenophonus lipoptenae (0.84 Mb), an obligate intracellular symbiont of the blood-feeding deer ked *L. cervi* (Nováková et al., 2016). Note that *Arsenophonus lipoptenae* was most closely related to *Arsenophonus* CJ based on the 16S rRNA sequences, as mentioned above (Figure 2B). In total, the *Arsenophonus* CJ genome encoded 512 proteins, 3 rRNAs (16S, 5S, and 23S), 32 tRNAs, and 11 pseudogenes. Notably, the *Arsenophonus* CJ genome exhibited a low coding density (60.3%), which may be a sign of ongoing genomic erosion.

The *Hamiltonella* CJ genome had a length of 2,258,324 bp, although we could not produce a circular contig (Figure 4C and Table 1). The genome size of *Hamiltonella* CJ was similar to that of *Hamiltonella* of *A. pisum* (Degnan et al., 2009), which is a facultative symbiont of aphids, and was larger than that of *Hamiltonella* of *Bemisia tabaci* (Rao et al., 2015), which is a co-obligate endosymbiont of whiteflies. The *Hamiltonella* CJ genome encoded 2,097 proteins, 9 rRNAs (3 of 16S, 3 of 5S, and 3 of 23S), 42 tRNAs, and 72 pseudogenes.

Based on two observations (i) the systematic co-infection of two endosymbionts, *Buchnera* and *Arsenophonus* in *Ce. japonica*, and (ii) the drastic genome reduction of these two bacteria, we hypothesized that the *Ce. japonica* aphid depends on *Buchnera* CJ and *Arsenophonus* CJ forming a co-obligate symbiosis. In several hemipteran insects, metabolic complementarity between co-symbionts has been observed (Husnik et al., 2013; Lamelas et al., 2011b; McCutcheon et al., 2009; Rao et al., 2015; Szabó et al., 2020; Wu et al., 2006). To determine if *Buchnera* CJ and *Arsenophonus* CJ form co-obligate associations with the host, we examined the genomic signature of metabolic complementarity by comparing the gene repertoires of these bacteria. We detected a complementarity gene repertoire was found in the metabolic pathways of riboflavin and peptidoglycan between *Buchnera* CJ and *Arsenophonus* CJ. While all genes responsible for riboflavin biosynthesis were missing from the *Buchnera* CJ genome, these genes were retained in the *Arsenophonus* CJ genome (Figure 4F and Table S4 and S5), indicating a complementary riboflavin-related gene set between the two symbionts. While *Buchnera* CJ lost all genes related to peptidoglycan synthesis (Table S4), the *Arsenophonus* CJ genome retained genes to synthesize dap-type peptidoglycan (*murABCDEFG*, *mraY*, *mrcB*, *ftsI*, and *dacA*) (Table S5). This complementary gene repertoire related to nutrition and cell wall components represents a genomic signature of co-obligate symbiosis.

Although *Arsenophonus*-related bacteria infect some aphids as facultative symbionts, the genus has not been reported as obligate symbionts in aphids, to the best of our knowledge. As such, we further inspected the genome of *Arsenophonus* CJ to understand the evolution of the obligate symbiosis. *Arsenophonus* CJ had a streamlined 853 Kb genome with an extremely low GC content (18.5%), a typical feature of obligatory endosymbionts. Notably, the *Arsenophonus* CJ genome exhibited a very low coding density (60.3%), suggesting ongoing genomic erosion. A COG analysis demonstrated that *Arsenophonus* CJ is missing many genes with broad functions, including essential housekeeping functions, such as “Replication, recombination and repair” (Figure 4E). For example, the *Arsenophonus* CJ genome lacked the SOS system genes *recA*, *lexA*, *umuCD,* and *uvrABC*, involved in DNA repair, as observed in *Buchnera* (Table S5). The *Arsenophonus* CJ genome lacked a majority of genes involved in amino acid biosynthesis pathways; however, it retained all genes responsible for riboflavin biosynthesis (Figure 4F and Table S5). Interestingly, genes for lipid A biosynthesis were missing or pseudogenized in the *Arsenophonus* CJ genome. The lipid A biosynthesis pathway is highly conserved in free-living bacteria and facultative bacterial symbionts; however, obligate intracellular symbionts often lack all or some components of this pathway, which is interpreted as an adaptation to the host immune responses (Wu et al., 2006). Among nine key genes involved in lipid A biosynthesis, five genes were lost and four were found to be pseudogenes in the *Arsenophonus* CJ genome (Table S7). The analysis of lipid A-related gene repertoire of the *Arsenophonus* CJ provided two insights: (1) *Arsenophonus* CJ lost the ability to produce functional lipid A, as observed in other obligatory symbionts, and (2) the degenerative process occurred relatively recently, since four pseudogenes were detected. Taken together, the *Arsenphonus* CJ genome showed the features of obligatory symbionts and the signature of genomic erosion due to the ongoing obligate interaction.

### Infection of *Buchnera*/*Arsenophonus* symbionts and host embryogenesis indicate developmental integration

Obligate symbionts often exhibit “developmental integration” with hosts, where the processes of symbiont infection and host oogenesis or embryogenesis are well-coordinated (Braendle et al., 2003; Miura et al., 2003; Shigenobu and Wilson, 2011). To address the developmental integration in the context of multi-partner symbiosis in *Ce. japonica*, we analyzed the formation of the bacteriome and dynamics of symbionts during parthenogenetic embryogenesis. We dissected ovarioles from the third or fourth instar nymphs of viviparous aphids, within which a series of developing embryos can be observed. Until anatrepsis (stage 8), when the germ band is invaginating from both the dorsal and ventral sides, no symbionts were detected (Figure 5A). Around the time of germ band elongation and folding into an S shape (stage 11), we found a mixed population of *Buchnera* and *Arsenophonus* incorporated into the embryo from the posterior part (Figure 5B). At the later stage of the germ band elongation, when it twisted with more abdominal segments (stage 12), we observed the first formation of bacteriocytes, where huge host nuclei and symbiont masses were cellularized into bacteriocytes (Figure 5C). At this stage, both *Buchnera* and *Arsenophonus* co-existed in the central bacteriocyte; however, only *Buchnera* cells were found in peripheral bacteriocytes (Figure 5C, 5D, and 5E). By the time of the initiation of limb bud formation (stage 13), several bacteriocytes aggregated into an organ-like structure (i.e., a bacteriome) located at the ventral part of the embryo. Within the bacteriome, *Buchnera* and *Arsenophonus* were segregated into different types of bacteriocytes: *Arsenophonus* resided in a single syncytial bacteriocyte, while *Buchnera* resided in multiple peripheral uninucleate bacteriocytes surrounding the *Arsenophonus*-containing bacteriocyte (Figure 5F). The central *Arsenophonus*-containing bacteriocyte was syncytial, containing four host nuclei, while the peripheral *Buchnera*-containing bacteriocyte was uninucleate. After katatrepsis, the bacteriome was located in the dorsal abdomen of the embryo (Figure 5G and 5H). At this stage, the morphology of the bacteriome changed from a round shape to a U-shape, maintaining the orientation of the two types of bacteriocytes—outer *Buchnera*-containing bacteriocytes and inner *Arsenophonus*-containing bacteriocytes. In the late embryo prior to larviposition, the bacteriome divided laterally to form a pair of bacteriomes at the anterior abdomen (Figure 5I). After larviposition, the pair of bacteriomes was located at the anterior abdomen, maintaining the orientation of the two types of bacteriocytes (Figure 6E). The morphology and localization of the bacteriomes in first instar larvae were comparable to those observed in adults (Figure 3A).

**Figure 5.**
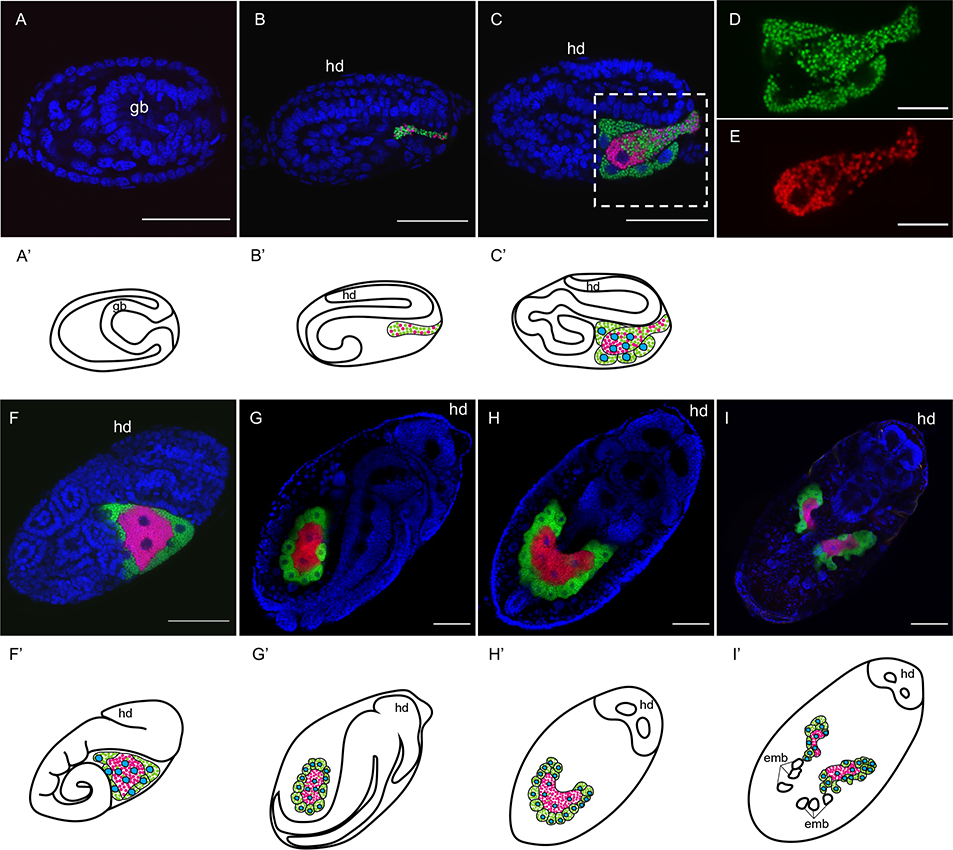
Infection and developmental integration of *Buchnera* and *Arsenophonus* symbionts during host embryogenesis. (A) Embryo during anatrepsis. Neither *Buchnera* nor *Arsenophonus* are observed at this stage. (B) S-shape embryo. Both *Buchnera* and *Arsenophonus* begin to infect the embryo from the posterior part. (C) Twisting embryo. Infection with both *Buchnera* and *Arsenophonus* is continuing. New bacteriocytes harboring *Buchnera* are formed. (D and E) Only Cy5 (D) and Cy3 (E) signals in C are shown. (F) Limb bud formation. Limb buds are formed in the thorax region and the germ band is elongating. Symbiont transmission is finished at this stage. (G and H) Germ band retraction is completed after katatrepsis. G and H show the lateral view and dorsal view of the same individual. The germ band is retracted to the posterior tip. One huge bacteriome exists in the abdomen. (I) An embryo prior to larviposition. The bacteriome is divided and forms a pair of bacteriomes. (A–I) Blue (DAPI), green (Cy5), and red (Cy3) indicate nuclei, *Buchnera*, and *Arsenophonus*, respectively. (A’–I’) Schematic diagram of images in (A–I). Cyan, green, and magenta indicate nuclei of bacteriocytes, *Buchnera*, and *Arsenophonus*, respectively. A–G: Lateral view, H and I: Dorsal view. Scale bar: 50 µm for A–H; 100 µm for I. gb, germ band; h, head.

**Figure 6.**
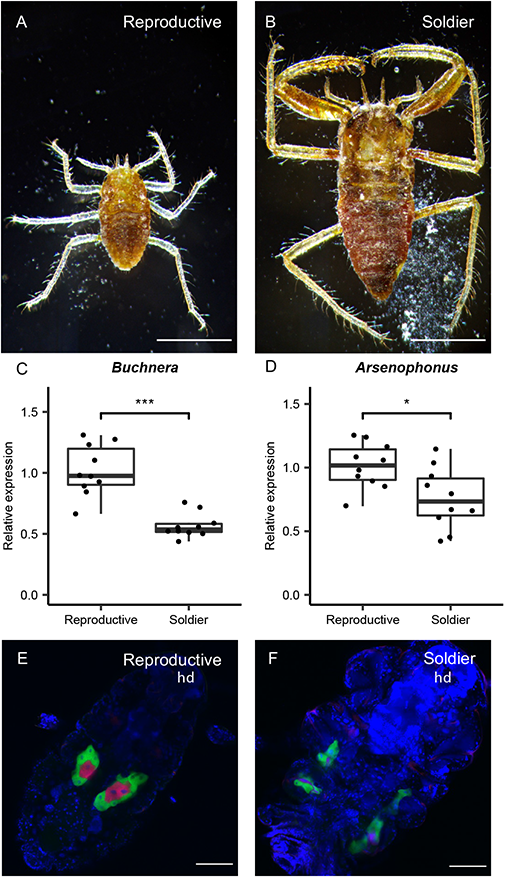
Comparison of symbiosis status between soldier and reproductive castes. (A and B) Light microscopic images of first-instar nymphs of reproductive (A) and soldier (B) castes. (C and D) Comparison of symbiont titers of first instar nymphs between castes. (C) *Buchnera* and (D) *Arsenophonus* titers. *Buchnera* and *Arsenophonus* were measured by qPCR using the bacterial symbiont *DnaK* gene standardized by the host *RpL7* gene. Asterisks indicate statistically significant differences (Welch’s *t*-test *P* < 0.001 in A, P < 0.05 in B). (E and F) Confocal images of the localization of both symbionts in the first instar nymphs of reproductive (E) and soldier (F) castes. Scale bars indicate 500 µm in (A and B) and 100 µm in (E and F). hd, head.

*Hamiltonella* CJ was also detected in the infecting bacterial mass together with *Buchnera* and *Arsenophonus* at the S-shape embryo stage (Figure S5A). However, unlike *Buchnera* and *Arsenophonus*, at later stages, *Hamiltonella* was scattered around the periphery of the bacteriome (Figure S5B) and was sometimes detected in or around unknown cells or organs, different from bacteriomes. Figure S5B shows an example of the localization near the tip of the germband.

### Symbiosis status differs between soldier and reproductive castes

*Ce. japonica* is eusocial and produces sterile individuals known as a soldier caste specialized for colony protection (Figure 1A, 6A, and 6B). With our laboratory rearing system, asexual viviparous females cultured on the bamboo *S. senanensis* produce the normal (reproductive) caste and the soldier caste. Soldiers remain first-instar nymphs throughout their life on the bamboo. We compared symbiosis status between castes. First, we evaluated *Buchnera* and *Arsenophonus* titers in first-instar individuals of the reproductive caste and soldier caste in *Ce. japonica* by quantitative PCR (Figure 6C and 6D). *Buchnera* was detected from both castes in *Ce. japonica*; however, the *Buchnera* titer was approximately 44% lower in the soldier caste than in the reproductive caste and the difference was statistically significant (Figure 6C; Welch’s *t*-test, p < 0.001). *Arsenophonus* was also detected from the both castes in *Ce. japonica*; however, the *Arsenophonus* titer was approximately 25% lower in the soldier caste than in the reproductive caste and this difference was statistically significant (Figure 6D; Welch’s *t*-test, p < 0.05). Second, in an analysis of morphological differences, found that soldiers possess a pair of bacteriomes with a different morphology from that in the reproductive caste (Figure 6E and 6F). Taken together, soldiers exhibit a different symbiotic condition distinct from that of the reproductive caste, including fewer symbionts in apparently malformed bacteriomes.

## Discussion

### *Arsenophonus* is an obligate symbiont in *Ce. japonica* forming dual symbiosis with *Buchnera*

We investigated bacterial symbionts of the eusocial aphid *Ce. japonica* by an integrative approach using molecular, genomic, and microscopic techniques. We found that two bacterial species, *Buchnera* CJ and *Arsenophonus* CJ, coexisted in all natural populations and individuals surveyed (Figure 1C, 1D, S1 and Table S1 and S2). The persistent presence of *Buchnera* in *Ce. japonica* is not surprising, as the species is a common obligatory endosymbiont in aphids (Baumann, 1995; Chong et al., 2019; Shigenobu and Wilson, 2011; Shigenobu et al., 2000). *Buchnera* CJ showed typical characteristics of *Buchnera* of other aphids, i.e., a small genome, transovarial inheritance, intracellular localization in bacteriocytes, and a subcellular morphology exhibiting a round-shaped bacterial cell encapsulated by the host-derived membrane (symbiosomal membrane). On the other hand, the systematic infection of *Arsenophonus* CJ was an unexpected and novel finding in this study. Although *Arsenophonus*-related bacteria infect some aphids as facultative symbionts (Ayoubi et al., 2020; Wagner et al., 2015; Wulff et al., 2013), the consistent infection of *Arsenophonus* in *Ce. japonica* suggests that the symbiosis is obligatory for the host. Microscopic observation showed that *Arsenophonus* cells were localized inside bacteriocytes, similar to the intracellular symbiosis observed in *Buchnera* (Figure 3E and 3G). Each *Arsenophonus* cell was surrounded by the host-derived membrane (symbiosomal membrane), as observed in *Buchnera* (Figure 3N). Furthermore, *Arsenophonus* symbionts were maternally and vertically inherited in the host offspring (Figure 5B–5I). *Arsenophonus* CJ had a small, streamlined genome (Figure 3B and Table 1), typical of obligate bacterial symbionts. Taken together, we detected a dual obligate symbiosis involving *Arsenophonus* sp. and *Buchnera* in *Ce. japonica*.

### Paraphyly of *Arsenophonus* genus and the origin of *Arsenophonus* CJ

The genus *Arsenophonus* is a cluster of bacterial symbionts found in taxonomically widespread insects (Nováková et al., 2009) and *Arsenophonus*-related bacteria have been reported to infect some aphids as facultative symbionts (Ayoubi et al., 2020; Jousselin et al., 2013; Wagner et al., 2015; Wulff et al., 2013). Our molecular phylogenetic analysis of 16S rRNA sequences indicated that *Arsenophonus* is a paraphyletic genus sharing a common ancestor with *Aschnera* and *Riesia*. Based on the phylogenetic tree, *Arsenophonus* and related bacteria could be clearly divided into two groups, Group A and Group B (Figure 2B). Group A mainly consisted of symbionts of blood-feeding insects, such as Hippoboscoidea flies and Anoplura sucking lice. Most *Arsenophonus*, *Aschnera,* and *Riesia* species included in Group A are obligate symbionts with small genomes ranging from 0.53 to 0.84 Mb (Boyd et al., 2017; Hosokawa et al., 2012; Nováková et al., 2016). For example, *Candidatus* Riesia pediculicola of primate lice, *Candidatus* Aschnera chinzeii of bat flies, and *Candidatus* Arsenophonus lipoptenae of deer ked *L. cervi* are obligate intracellular symbionts of hematophagous insects. In all cases, these endosymbionts reside within bacteriocytes and supply hosts with essential B vitamins that are deficient in the blood diet (Boyd et al., 2017; Hosokawa et al., 2012; Nováková et al., 2016; Sasaki-Fukatsu et al., 2006). Group B was mainly composed of symbionts of hemipteran insects, including aphids (Figure 2B). *Arsenophonus* species of Group B had relatively large genomes and appeared to be facultative symbionts, such as the son-killer bacterium *Arsenophonus nasoniae* (4.99 Mb) of *Nasonia vitripennis* (Darby et al., 2010), *Arsenophonus triatominarum* (3.86 Mb) of *Triatoma infestans*, and *Arsenophonus* sp. (2.42 Mb) of *Aphis craccivora*, except an obligate endosymbiont of a whitefly, *Aleurodicus disperses* (0.66 Mb) (Santos-Garcia et al., 2018). In sum, *Arsenophonus* species can be classified into two distinct groups, a group of obligatory endosymbionts of hematophagous insects and a group of facultative symbionts of Hemiptera. It should be noted that *Arsenophonus* CJ belonged to Group A, supporting the obligate nature of the observed symbiosis.

The phylogenetic position of *Arsenophonus* CJ raises a question about its origin. *Arsenophonus* CJ was assigned to Group A (i.e., hematophagous insect endosymbionts), suggesting that it originated from symbionts of hematophagous insects by horizontal transfer. Alternatively, since *Arsenophonus* is widespread in Hormaphidinae aphids (Xu et al., 2021), the ancient acquisition of this symbiont by *Ce. japonica* is also possible. However, it is clear that *Arsenophonus* CJ evolved from ancestors distinct from Group B. Extensive phylogenetic analyses of *Arsenophonus* in Hormaphidinae aphids with more extensive taxon sampling are needed to understand the origin of co-obligate *Arsenophonus* symbionts.

### *Buchnera* and *Arsenophonus* as co-obligate symbionts

Our genome analysis revealed that the genome size of *Buchnera* CJ (Table 1) was significantly smaller than the majority of *Buchnera* genomes (Chong et al., 2019; Shigenobu and Yorimoto, 2022). It is possible that a loss of some functions in this *Buchnera* lineage was compensated by the other obligate symbiont *Arsenophonus* CJ. We examined the genomic signature of metabolic complementarity. All genes responsible for riboflavin biosynthesis were missing from the *Buchnera* CJ genome but present in the *Arsenophonus* CJ genome (Figure 4F and Table S4 and S5), indicating a complementary riboflavin-related gene set between two symbionts. *Arsenophonus* CJ may provide riboflavin to the host, instead of *Buchnera*, which provides riboflavin in the Aphid– *Buchnera* symbiosis (Nakabachi and Ishikawa, 1999). Similarly, peptidoglycan synthesis genes were lost in *Buchnera* CJ (Table S4) and present in the *Arsenophonus* CJ genome (particularly *murABCDEFG*, *mraY*, *mrcB*, *ftsI*, and *dacA*) (Table S5). These results provide evidence for complementary repertoires of genes involved in nutrition and cell wall production in *Buchnera* CJ and *Arsenophonus* CJ, further indicating that they represent a co-obligate symbiosis.

Our histological observation revealed a characteristic arrangement of the two symbionts within the same bacteriome. Both symbionts resided within the same bacteriome but in separate bacteriocytes; the central syncytial bacteriocytes harboring *Arsenophonus* CJ were morphologically different from the peripheral uninucleate bacteriocytes harboring *Buchnera* within the bacteriome of *Ce. japonica* (Figure 3). In bacteriome formation during embryogenesis, although both *Buchnera* CJ and *Arsenophonus* CJ were maternally transmitted simultaneously to the embryos, the symbionts were segregated into different types of bacteriocytes (Figure 5B–F). These observations indicate a high level of anatomical and developmental integration of the two symbionts and the host.

### Parallel evolution of co-obligate symbiosis in aphids

The co-obligate symbiosis of *Buchnera* and *Serratia* has occurred in aphid lineages, such as *Cinara* and *Tuberolachnus* in the subfamily Lachninae, *Periphyllus* in the subfamily Chaitophorinae, and *Aphis urticata* and *Microlophium carnosum* in the subfamily Aphidinae (Lamelas et al., 2011a, 2011b; Manzano-Marín and Latorre, 2014; Manzano-Marín et al., 2016; Monnin et al., 2020; Pérez-Brocal et al., 2006). In several *Cinara* species, co-obligate *Serratia* has been replaced by *Erwinia*, *Fukatsuia*, and *Sodalis* symbionts (Manzano-Marín et al., 2017; Manzano-Marín et al., 2019; Meseguer et al., 2017). Our study is the first report of co-obligate symbiosis in the subfamily Hormaphidinae and revealed a novel partner combination (i.e., *Buchnera* and *Arsenophonus*). Regardless of the aphid lineage or co-obligate symbiont species, there are amazing similarities among co-obligate symbioses ranging from genomic features to the morphology of bacteriocytes. While typical *Buchnera* genome sizes are ∼0.6 Mb, those of co-obligate *Buchnera* in *Ce. japonica*, Lachninae, and *Periphyllus* aphids have ∼0.4 Mb genomes, regardless of phylogenetic positions (Table 1) (Chong et al., 2019; Shigenobu and Yorimoto, 2022). All of these ∼0.4 Mb *Buchnera* genomes lost genes in the riboflavin, ornithine, and peptidoglycan biosynthesis pathways, although the ornithine and peptidoglycan biosynthesis pathways were also lost in several *Buchnera* genomes of other subfamilies (Figure 4F and Table S4) (Chong et al., 2019; Lamelas et al., 2011a; Manzano-Marín et al., 2016; Manzano-Marín et al., 2019; Monnin et al., 2020; Pérez-Brocal et al., 2006). The loss of functions in *Buchnera* is presumably compensated by the partner obligate symbiont, as evidenced by the complementary gene repertoire. In dual symbiosis, *Buchnera* and co-infected symbionts, such as *Serratia*, *Erwinia*, *Fukatsuia*, or *Sodalis*, are found in the same bacteriome but sorted separately into the distinct bacteriocytes in Lachninae aphids (Manzano-Marín et al., 2017; Manzano-Marín et al., 2019; Meseguer et al., 2017). Mitochondrial density differs between *Buchnera*-containing bacteriocytes and co-obligate symbiont-containing bacteriocytes: mitochondrial abundance was lower in bacteriocytes harboring *Serratia* in *Ci. cedri* (Soriano-Navarro et al., 2004) and *Arsenophonus* in *Ce. japonica* (Figure 3J and 3K). These common characteristics are consistent with parallel evolution of the co-obligate symbiosis in aphid lineages, implying that there are evolutionary constraints on the aphid–*Buchnera* symbiosis.

### Roles of Arsenophonus CJ

Based on the genomic analysis, *Arsenophonus* CJ is likely involved in riboflavin provisioning to the host (Figure 4F). A similar function in B-vitamin provisioning has been reported for whiteflies, lice, and Hippoboscoidea flies (Nováková et al., 2015, 2016; Santos-Garcia et al., 2018). An experimental study has demonstrated that a facultative *Arsenophonus* symbiont expands the dietary breadth in the aphid *Aphis craccivora* (Wagner et al., 2015). *Arsenophonus* CJ may also contribute to the usage of host plants, the snowbell tree *Styrax japonicus* and bamboo grasses, such as *Pleioblastus chino*, *P. simonii* and *Sasa senanensis*. However, the acquisition of new co-obligate symbiont species is not associated with adaptation to novel ecological niches in the evolution of co-obligate symbiosis of *Cinara* aphids (Meseguer et al., 2017). Further studies are needed to clarify the roles of the *Arsenophonus* symbiont in *Ce. japonica*.

### Ongoing reductive evolution of the *Arsenophonus* CJ genome

Synteny analysis showed that *Arsenophonus* genomes have experienced many rearrangements and deletions (Figure S4C and S4D). In addition, the *Arsenophonus* CJ genome showed a low coding density and multiple pseudogenes (Table 1). Recently evolved symbionts are characterized by a low coding density, pseudogene formation, and many genome rearrangements and deletions (Burke and Moran, 2011; Ochman and Davalos, 2006). Accordingly, *Arsenophonus* CJ is a recently evolved co-obligate symbiont and gene inactivation is ongoing. An interesting example of ongoing genomic erosion in *Arsenophonus* CJ was the gene repertoire for lipid A biosynthesis. Lipid A is the hydrophobic anchor of lipopolysaccharide in the outer membrane of gram-negative bacteria (Raetz and Whitfield, 2002; Raetz et al., 2009), and elicits a strong immune response in host animals (Raetz and Whitfield, 2002). The key genes for lipid A biosynthesis were either missing or pseudogenes in the *Arsenophonus* CJ genome (Table S7). Losing the ability to produce lipid A in *Arsenophonus* CJ may be beneficial for the symbiotic association with host animals by accommodating the host immune response. The observed pseudogenization suggests an ongoing genomic erosion of *Arsenophonus* CJ, probably leading to a more streamlined obligatory endosymbiont specialized for the *Arsenophonus*–*Buchnera* dual symbiosis.

### Symbiosis in the sterile soldier in eusocial aphids

Eusocial aphid species have been found in only two subfamilies, Hormaphidinae and Eriosomatinae (Pike and Foster, 2008; Stern and Foster, 1996). The eusocial aphid *Ce. japonica* provides a unique opportunity to study how symbiosis operates in a caste system. In a comparison of normal (reproductive) nymphs and sterile soldiers, we detected significantly fewer symbionts in soldiers than in normal nymphs (Figure 6C and 6D) as well as a distorted bacteriome shape (Figure 6E and 6F), indicating that the symbiotic condition differs between social castes in *Ce. japonica*. Soldiers of eusocial aphids are sterile and do not grow after birth (Chung and Shigenobu, 2022; Stern and Foster, 1996). Such differences in nutritional conditions between castes may be related to the difference in symbiotic status. In the eusocial aphid *Colophina arma* of the subfamily Eriosomatinae, the soldier caste lacks endosymbionts and bacteriomes entirely (Fukatsu and Ishikawa, 1992). In the eusocial ant tribe Camponotini, the obligate symbiont *Blochmannia* enhances nutrition, thereby influencing the size of worker ants (Feldhaar et al., 2007; Sinotte et al., 2018; Zientz et al., 2006). The mechanisms and biological significance of caste-dependent symbiosis in social insects are intriguing subjects for future exploration.

### Limitations of the study

Our study found a dual obligate symbiosis involving *Arsenophonus* sp. and *Buchnera* in *Ce. japonica* with molecular, genomic, and microscopic approaches. Genomic analysis showed that *Arsenophonus* CJ likely has a role involved in riboflavin provisioning to the host. However, genomic data cannot provide direct evidence of whether riboflavin provisioning by *Arsenophonus* CJ is essential for the hosts and restrict speculation of other roles of the symbionts. Antibiotic treatment to eliminate each symbiont specifically is needed to solve these problems. We revealed streamlined small genomes and phylogenetic positions of *Buchnera* and *Arsenophonus* of *Ce. japonica*. Surprisingly, molecular phylogenetic analysis based on the 16S rRNA sequences showed that *Arsenophonus* of *Ce. japonica* was related to endosymbionts of hematophagous Diptera (the superfamily Hippoboscoidea), not included in the clade of *Arsenophonus* symbionts in Hemiptera. To understand the origin of *Arsenophonus* CJ and the timing of drastic genome reduction in the *Buchnera* and *Arsenophonus*, genome sequencing of bacterial symbionts with more extensive taxon sampling in the subfamily Hormaphidinae is needed.

## Acknowledgments

We appreciate Dr. Katsushi Yamaguchi and Dr. Asaka Akita in the Functional Genomics Facility, NIBB Core Research Facilities for technical support with Illumina library preparation and sequencing. We appreciate Dr. Yuuki Kobayashi, Dr. Wen Hsin-i, and Mika Ikeda for the technical support with Nanopore library preparation and sequencing. We appreciate Dr. Chen-yo Chung, Yasuhiko Chikami, Dr. Naoki Minamino, and Dr. Yasuhiro Kamei for technical advices. We appreciate Dr. Shinichi Yoda, Yasuhiko Chikami, and Kathrine Tan for support and discussion. Computational resources were provided by the Data Integration and Analysis Facility, National Institute for Basic Biology. This work was funded by a Grant-in-Aid for Scientific Research (B) (no. 17H03717) to S. Shigenobu, Grant-in-Aid for Scientific Research (A) (no. 20H00478) to S.S., Grant-in-Aid for Scientific Research on Innovative Areas (Research in a Proposed Research Area) (no. 17H06384) to Dr. Shigeru Kuratani and Grant-in-Aid for JSPS Fellows (no. 20J11981) to S. Yorimoto.

## Author contributions

S. Y. and S. S. designed the research. S. Y. and M. H. collected aphid samples. S. Y. and M. K. performed TEM. S. Y. performed molecular biology experiments and microscopic analyses. S. Y. and S. S. performed next-generation sequencing and bioinformatics analyses. S. Y. and S. S. wrote the manuscript. All authors provided comments on revisions and approved the final manuscript.

## STAR Methods text

### RESOURCE AVAILABILITY

#### Lead contact

Further information and requests for resources and reagents should be directed to and will be fulfilled by the lead contact, Dr. Shuji Shigenobu.

#### Materials availability

The isofemale NOSY1 strain of *Ce. japonica* established in this study has been maintained at the Laboratory of Evolutionary Genomics, National Institute of Basic Biology.

#### Data and code availability

- All sequencing data and assembled genome data have been deposited at the DDBJ under the bioProject ID: PRJDB13695. Accession numbers are listed in the Key Resources Table.
- Original codes are available on GitHub (https://github.com/shigenobulab/21shunta_cjsym_msprep). Annotation data are available through Figshare (DOI: 10.6084/m9.figshare.c.6026333).
- Any additional information required to reanalyze the data reported in this paper is available from the lead contact upon request.

### EXPERIMENTAL MODEL AND SUBJECT DETAILS

#### Aphid rearing

An isofemale strain of the aphid *Ce. japonica*, strain NOSY1, was established from a single female collected from a secondary host plant, bamboo grass *Sasa senanensis*, at the foot of Mt. Norikura, Nagano Prefecture, Japan (1525.7 m above the sea level; 36°07’09.0″N, 137°37’18.4″E). The strain has been maintained in the asexual viviparous phase on *S. senanensis* at 20 °C with a long-day condition (16 hours light/8 hours dark cycle) and 60% relative humidity in our laboratory since September 2017. *S. senanensis* was grown singly in 0.6L pots filled with potting soil at 25 °C before use for feeding for aphids. For the species identification, a partial cytochrome c oxidase I (COI) sequence (658 bp) of *Ce. japonica* registered in NCBI (EU701571.1) and the mitochondrial genome sequence obtained from our *de novo* assembly data of *Ce. japonica* strain NOSY1 (see “Symbiont genome assembly and annotation” section) were aligned using MAFFT (v7.490) (Katoh, 2002) with the default options, which resulted in only one nucleotide mismatch.

### METHOD DETAILS

#### Aphid collection

Twenty-six colonies of *Ceratovacuna japonica* (Takahashi, 1958) were collected at nine geographically distinct populations from three different host plant species across Japan from 2006 to 2019. The host plants and locations of sampling are listed in Table S1 and S2. These aphid samples were preserved in 100% ethanol until 16S ribosomal DNA amplicon sequencing was performed.

#### 16S ribosomal DNA amplicon sequencing analysis

The genome DNA (gDNA) of *Ce. japonica* colonies was extracted using DNeasy Blood & Tissue Kit (Qiagen, Hilden, Germany) according to the manufacturer’s instruction. Aphids were rinsed with fresh 70% ethanol with vigorous agitation to remove wax on the surface of their bodies. Each gDNA sample was extracted from 6 to 20 individuals of the reproductive caste. gDNA samples with low concentration were concentrated by ethanol precipitation with Ethachinmate (Nippon Gene, Tokyo, Japan). The 16S rDNA libraries were prepared using Quick-16S NGS Library Prep Kit (Zymo Research, Irvine, CA, USA) targeting hypervariable 16S rRNA regions, V1–V2 and V3–V4 regions, with one negative (ZymoBIOMICS^®^ DNase/RNase Free Distilled Water, Zymo Research) and one positive control (ZymoBIOMICS^®^ Microbial Community DNA Standard, Zymo Research). Concentrations of the libraries were quantified with KAPA SYBR FAST qPCR Kit (Kapa Biosystems, Wilmington, MA, USA) by Applied Biosystems 7500 Real-Time PCR Systems (Applied Biosystems, Foster City, CA, USA). The libraries were sequenced using the Illumina MiSeq platform (Illumina, Foster City, CA, USA), and 250 bp of paired-end reads were generated. The sequencing of 16S rRNA hypervariable regions, V1–V2 and V3–V4, yielded 360,704 and 484,288 raw reads, respectively. The Illumina raw reads were deposited in the DDBJ DRA database under accession number DRA014323.

Raw paired-end reads were analyzed using QIIME 2 (version 2020.8) (Bolyen et al., 2019) with the following plugins: dada2 (Callahan et al., 2016) for quality filtering, trimming length, merge paired reads and removing chimeric sequences; naïve Bayes classifier (Bokulich et al., 2018) for taxonomy assignment against the Silva132 database (Quast et al., 2013). Forward and reverse primer regions were trimmed from the raw reads as follows: 19 bp from 5’ end of forward reads and 16 bp from 5’ end of reverse reads in the V1–V2 amplified data, 16 bp from 5’ end of forward reads and 24 bp from 5’ end of reverse reads in the V3–V4 amplified data. After quality filtering and removing chimeric sequences, 332,656 and 432,594 reads from the V1–V2 and V3–V4 regions were remained, respectively.

#### Diagnostic PCR

Genomic DNA was prepared from *Ce. japonica* colonies in the same way as described in the “16S ribosomal DNA amplicon sequencing analysis” section. Same gDNA samples corresponding to natural populations of *Ce. japonica* (#1–#26) were used to confirm the presence or absence of symbionts. In addition, gDNA from each of the 12 adult individuals (populations of #2–4, #14, #18–20, and #24–26) was used to confirm the infection status at the individual level. Primers targeting a single-copy gene, *dnaK*, for *Buchnera*, *Arsenophonus*, and *Hamiltonella* were designed using Primer3Plus (Untergasser et al., 2007): CjBucDnaK_FW2 (5’-CAG CAG ATT CAT CTG GAC CTA AAC -3’) and CjBucDnaK_RV2 (5’-CCA TAG GCA TTC TAG TTT GAC CAC -3’) primers for *Buchnera*; CjArsDnaK_FW2 (5’-TGG AAT TCA AGC AGC ACC AC -3’) and CjArsDnaK_RV2 (5’-TCT GCA TTT GCT TCT GCA TC -3’) primers for *Arsenophonus*; CjHamDnaK_F3 (5’-ATG CAC TGA CGA TGG TTT CTG C -3’) and CjHamDnaK_R3 (5’-ACT CAG CAT CAA CAG CAT CTG C -3’) for *Hamiltonella*. PCR reaction mixtures were comprised of 10 µl KOD SYBR^®^ qPCR Mix (Toyobo, Osaka, Japan), 0.6 µl of each primer (0.3 µM), 2 µl gDNA, and 6.8 µl UltraPure™ DNase/RNase-Free Distilled Water (Invitrogen, Carlsbad, CA, USA). PCR was performed on LightCycler^®^ 96 (Roche, Basel, Switzerland) with the following program: 98°C for 2 min, followed by 30 cycles consisting of 98°C for 10 sec, 60°C for 10 sec, and 68°C for 30 sec. The PCR products and FastGene 100 bp DNA Marker (Nippon Genetics) were loaded on 1.8% UltraPure^TM^ Agarose (Invitrogen) gels containing SYBR Safe DNA Gel Stain (Invitrogen) with 100V for 25 min and imaged on the Molecular Imager^®^ ChemiDoc^TM^ XRS+ system using UV transillumination (Bio-Rad, Hercules, CA, USA).

#### Real-time quantitative PCR

Genome DNA was extracted from first-instar nymphs of the reproductive caste and soldier caste in the strain NOSY1 in the same way as described in the “16S ribosomal DNA amplicon sequencing analysis” section. Each sample contained the gDNA of 3 individuals. We used the primers targeting the *dnaK* gene: CjBucDnaK_FW2 and CjBucDnaK_RV2 for *Buchnera* and CjArsDnaK_FW2 and CjArsDnaK_RV2 for *Arsenophonus*. We additionally designed primers targeting the *RpL7* gene in the host *Ce. japonica* as an internal control: Cjap_TR33288_RpL7_F1 (5’-GGC CTT TCA AAT TAA ACA CCC CAA C -3’) and Cjap_TR33288_RpL7_R1 (5’-ATC TTC CCG GTT TCC AAA GTC G -3’) primers. PCR reaction mixtures were composed of 10 µl KOD SYBR^®^ qPCR Mix (Toyobo), 0.6 µl of each primer (0.3 µM), 2 µl gDNA, and 6.8 µl UltraPure™ DNase/RNase-Free Distilled Water (Invitrogen). Real-time quantitative PCR was performed in 40 cycles using LightCycler^®^ 96 Instrument (Roche) with the following program: 98°C for 2 min, followed by 40 cycles consisting of 98°C for 10 sec, 60°C for 10 sec, and 68°C for 30 sec. The amount of the *dnaK* genes of bacterial symbionts was normalized by the *RpL7* gene of the host and the relative amount was calculated using the ΔΔCt methodology (Livak and Schmittgen, 2001).

#### Hologenome sequencing

Nanopore long reads and Illumina short reads were generated to achieve high-quality symbiont genome assemblies. For Nanopore library preparation, 19.69 mg of 25 adult individuals of the strain NOSY1 was used. Aphids were rinsed with fresh 70% ethanol with vigorous agitation to remove wax on the surface of their bodies. To prepare the high molecular weight (HMV) DNA of aphids, frozen aphids were transferred to a mortar and gently ground into a fine powder with liquid nitrogen. Frozen powdery QIAGEN G2 buffer (Qiagen), which was generated by adding 2-mercaptoethanol to QIAGEN G2 buffer and spraying the mixed buffer into liquid nitrogen in a glass beaker, was added to the sample and blended quickly. Letting the mixture thaw in a tube, RNaseA (Qiagen) and Proteinase K (Qiagen) were added, and the sample was incubated at 40°C for 3.0 hours without agitation. The sample was centrifuged at 9,500 rpm at 4 °C for 20 minutes and the supernatant was subjected to DNA extraction with a QIAGEN Genomic-tip 20/G column. The gDNA was eluted with 800 µl of Buffer QF twice, a 0.7-fold volume of isopropanol was added, and then the gDNA was centrifuged at 9,500 rpm at 4°C for 20 minutes. The pellet was washed twice using fresh 70% ethanol and centrifuged at 15,000 rpm at 4°C for 5 minutes. The pellet was dried for a few minutes and then the gDNA was eluted with 51 µl TE buffer at room temperature overnights. Note that the extracted gDNA includes genomes derived from aphids and the symbionts, i.e., hologenome. The quantity of extracted gDNA was measured using Qubit dsDNA HS Assay Kit (Thermo Fisher Scientific, Waltham, MA, USA) and Qubit^®^ 2.0 Fluorometer (Thermo Fisher Scientific). The quality of extracted gDNA was assessed using Nanodrop ND-2000C (Thermo Fisher Scientific). The integrity of the HMW genome was assessed by a pulsed-field gel electrophoresis using CHEF Mapper (Bio-Rad). With this HMW genome, a Nanopore sequencing library was prepared using the SQL-LSK110 Ligation Sequencing Kit (Oxford Nanopore Technologies, Oxford, UK) according to the manufacturer’s instruction and sequenced using the R10.3 flow cell on the GridION system. Reads were basecalled using GUPPY (version 4.3.4). The total number of raw Nanopore reads was 646,913. The raw Nanopore reads were deposited in the DDBJ database under accession number DRR379965.

For Illumina library preparation, 10.93 mg of 15 adult individuals of the strain NOSY1 was used. Aphids were rinsed with fresh 70% ethanol with vigorous agitation to remove wax on the surface of their bodies. Genomic DNA was extracted using DNeasy Blood & Tissue Kit (Qiagen) and then the gDNA was purified using the Genomic DNA Clean & Concentrator Kit (Zymo Research) according to the manufacturer’s instruction. The gDNA was fragmented into 200-500 bp (peak at 350 bp) using Covaris Focused-ultrasonicator M220. A library for whole genome sequencing was prepared using the TruSeq DNA PCR-Free Library Prep Kit (Illumina) according to the manufacturer’s instruction. Quality of the library was validated by the TapeStation HS D5000 (Agilent Technologies, Santa Clara, CA, USA). A concentration of the library was quantified by Applied Biosystems 7500 Real-Time PCR Systems (Applied Biosystems). The Illumina library was sequenced using the Illumina HiSeq X Ten platform (Illumina) at Macrogen Japan (Tokyo, Japan) with the 2x 150 bp paired-end sequencing protocol. The total number of raw Illumina paired-end reads was 204,892,576. The raw Illumina reads were deposited in the DDBJ database under accession number DRR379966.

#### Genome assembly and annotation

Raw Nanopore reads obtained from hologenomic samples (see above the section “Hologenome sequencing”) were used to assemble symbiont genomes and a mitochondrial genome of the host using raven (version 1.5.0) (Vaser and Šikić, 2021). Plasmid sequences of *Buchnera* were searched from the raw Nanopore reads using BLASTN (version 2.12.0) (Altschul et al., 1990). The raw Nanopore reads were mapped to the plasmid sequences using minimap2 (version 2.17-r941) (Li, 2018), and the sequences were polished once using Racon (version 1.4.20) (Vaser et al., 2017). Then, assembled symbiont genomes and the plasmids are polished using medaka (version 1.4.1, https://github.com/nanoporetech/medaka). Adapter trimming and quality filtering were performed on raw Illumina paired-reads using cutadapt (version 2.10) (Martin, 2011) and 182,821,759 (89.2%) paired-reads were passed. The cleaned Illumina reads were mapped to the assembled genomes and the plasmids using Bowtie2 (version 2.4.2) (Langmead and Salzberg, 2012) and then Pilon (version 1.24) (Walker et al., 2014) was used for assembly polishing. This polishing step with Bowtie2 and Pilon was repeated three times. In *de novo* genome assembling, 17,120 (read depth: 126.6 x), 13,819 (39.7 x), 44970 (97.3 x), and 10,873 (813.4 x) of 646,913 raw Nanopore long reads were used for the *Buchnera*, *Arsenophonus*, *Hamiltonella*, and mitochondrial genomes, respectively. For polishing the plasmid sequences with Racon, 335 (83.9 x) and 668 (130.6 x) of 646,913 raw Nanopore reads were used for pLeu and pTrp, respectively. For polishing the genome assembly with Illumina reads with pilon, 5,401,205 (1,819.4 x), 2,508,501 (413.9 x), 11,614,114 (702.9 x), 1,223,783 (8,126.6x), 180,211 (1,882.0 x), and 304,557 (2,228.9 x) of 182,821,759 cleaned Illumina reads were used for the genomes of *Buchnera*, *Arsenophonus*, *Hamiltonella*, and mitochondrion, and the plasmids of pLeu and pTrp, respectively. The assembled genome and the plasmid sequences were deposited in the DDBJ database under accession numbers: AP026065–AP026067 for *Buchnera*, AP026064 for *Arsenophonus*, and AP026068 for *Hamiltonella*.

Gene predictions were performed using Prokka (version 1.14.6) (Seemann, 2014). Pseudogene candidates were predicted using Pseudofinder (version 1.0) and determined these as pseudogenes according to the following criteria: genes less than 70% of the average length of DIAMOND BlastP hits and/or interrupted by pre-mature terminal codons. Functional annotations were performed using eggNOG-mapper (version 2.1.3) (Huerta-Cepas et al., 2017, 2019), the annotations include Cluster of Orthologous Genes (COG) category tags (Tatusov et al., 2000) and Kyoto Encyclopedia of Genes and Genomes (KEGG) IDs (Kanehisa and Goto, 2000). Metabolic pathways were reconstructed using the KEGG mapper tool (Kanehisa and Sato, 2020). Three symbiont genomes and two plasmids were visualized with annotations using circlize (version 0.4.15) (Gu et al., 2014) and gggenes (version 0.4.1) (Wilkins and Kurtz, 2019), respectively. To observe genome synteny, symbiont chromosomes are compared using BLASTN with dc-megablast task option (Altschul et al., 1990). The gene annotation files and scripts are available on Figshare (DOI: 10.6084/m9.figshare.c.6026333) and GitHub (https://github.com/shigenobulab/21shunta_cjsym_msprep), respectively.

#### Molecular phylogenetic analysis

To reconstruct phylogenetic trees with symbiont sequences of Ce. japonica obtained in this study, 16S rRNA sequences of relatives were downloaded from NCBI. See Table S3 for the accession number of the 16S rRNA sequences used for phylogenetic analyses. Multiple alignments of 16S rRNA sequences were performed using MAFFT (version 7.490) (Katoh, 2002). Gaps in the alignments were stripped using TrimAL (version 1.4.rev15) with “-gt 0.9” option (Capella-Gutiérrez et al., 2009). To find the best models for phylogenetic analysis, ModelTest-NG (version 0.1.6) (Darriba et al., 2020) was used; all of the bayesian information criterion, Akaike information criterion, and Akaike information criterion corrected for small samples supported consistently the GTR+I+G4 model and TVM+I+G4 model for *Buchnera* and *Arsenophonus*, respectively, for the Maximum likelihood (ML) trees. ML trees were reconstructed using RAxML-NG (version 1.0.3) (Kozlov et al., 2019) with the aforementioned models. Bootstrap values for ML phylogeny were obtained by 1,000 replicates. Phylogenetic trees were visualized using FigTree (version 1.4.4) (http://tree.bio.ed.ac.uk/software/figtree/). A multiple alignment of *Arsenophonus* of *Ce. japonica*, *L. cervi*, and *T. parasiticus* and a pairwise alignment of *Hamiltonella* of *Ce. japonica* and *A. pisum* were performed using MAFFT (v7.490) and visualized using Jalview (version 2.11.0) (Waterhouse et al., 2009).

#### Histology

Heads, tails, and appendages of adult insects were removed in a Bouin’s fixative (saturated picric acid: formaldehyde: glacial acetic acid = 15:5:1) to facilitate infiltration of reagents into the insect tissues at room temperature. The insects were fixed in fresh Bouin’s fixative at 4°C overnight, washed with 90% ethanol added lithium carbonate, and preserved in 90% ethanol at 4°C until use. Subsequently, the insects were dehydrated and cleared through an ethanol-butanol series. The cleared insects were immersed and embedded in paraffin at 60°C. The paraffin blocks were polymerized at 4°C and cut into 5 µm thick sections using the rotary microtome Leica RM 2155 (Leica, Wetzlar, Germany). The sections were mounted on microscopic slides coated with egg white-glycerin and deparaffinized by Lemosol A (FUJIFILM Wako Pure Chemical Corporation, Osaka, Japan). The deparaffinized sections were hydrated through an ethanol-water series and stained with Delafield’s hematoxylin. The stained sections were washed with 1% hydrochloric acid-ethanol for 40 seconds and then stained with eosin. The stained sections were dehydrated through a water-ethanol series and cleaned by Lemosol A. The sections were mounted into Canada balsam and the images were taken using the digital camera DS-FIL-U2 (Nikon, Tokyo, Japan).

#### Fluorescence *in situ* hybridization

Adults and first-instar nymphs were fixed by Carnoy’s fixative (ethanol: chloroform: acetic acid = 6:3:1) overnight and decolorized by the alcoholic 6% H_2_O_2_ solution for several weeks at room temperature. Late embryos were fixed by Carnoy’s fixative for 30 minutes and decolorized by the alcoholic 6% H_2_O_2_ solution overnight. H_2_O_2_ solution was exchanged every two or three days during incubation. Young embryos and dissected bacteriomes were fixed by 4% paraformaldehyde dissolved 1x phosphate buffered saline (PBS) (pH 7.4). The fixed samples were treated with 100% methanol for 30 minutes except for bacteriome samples. The samples were hydrated in a graded series of 0.3% Triton X-100 in 1x PBS prior to the hybridization. Hybridization was performed in 20 mM Tris-HCl (pH 8.0), 0.9 M NaCl, 0.01% SDS, 30% (vol/vol) formamide containing 100 nM each of fluorescent-labeled oligonucleotide DNA probes: Cy5_CjapBuc16S_1 (5’ -Cy5-CCT CTT CTA AGT AGA TCC-3’), Cy3-CjapArs16S_2 (5’-Cy3-CCC GAC CGA ATC GAT GGC-3’) and Cy5_CjapHam16S_V1V2 (5’ -Cy5-CTC AGT AAA CTG CGC TCA C-3’), which are complementary to 16S rRNA of *Buchnera*, *Arsenophonus,* and *Hamiltonella*, respectively, at room temperature for 2 hours. Excess probes were washed out with 0.3% Triton X-100 in 1x PBS. DNA was counterstained with DAPI (Dojindo Laboratories, Kumamoto, Japan) or Hoechst 33342 (Thermo Fisher Scientific). F-actin was stained with Alexa Fluor™ 594 phalloidin (Thermo Fisher Scientific) in 1x PBS at room temperature for 2 hours. Membranes were stained with FM™ 4-64FX (Thermo Fisher Scientific) in 1x PBS for 10 minutes on ice; then fixed by 4% paraformaldehyde dissolved 1x PBS for one hour on ice. The specimens were mounted with VECTASHIELD (Vector Laboratories, Burlingame, CA, USA) and analyzed under Olympus FLUOVIEW FV1000 confocal laser scanning microscope (Olympus, Tokyo, Japan). The sizes of bacteriomes were measured using FIJI (Schindelin et al., 2012).

#### Transmission electron microscopy (TEM)

Bacteriomes were dissected from adult aphids in 2.5% glutaraldehyde in 1x PBS (pH 7.4). The fixed samples were then postfixed with 1% osmium tetroxide for 1 hour. After washing with the same buffer, the specimens were dehydrated in a graded ethanol series at room temperature. The samples were treated with propylene oxide and infiltrated with propylene oxide-Epon (Epon 812 resin; TAAB Laboratories, Aldermaston, UK) solution (propylene oxide-Epon resin, 1:1 [v/v]) overnight. The samples were then embedded in Epon resin that was allowed to polymerize at 60°C for 72 hr. Ultrathin sections were cut on an ultramicrotome (Leica) and mounted on nickel grids. The sections were then stained with 4% uranyl acetate and lead citrate. After staining, all sections were examined under a transmission electron microscope (model JEM1010; JEOL, Tokyo, Japan) operated at 80 kV.

## QUANTIFICATION AND STATISTICAL ANALYSIS

The symbiont titers between castes were evaluated using Welch’s t-test on the ΔΔCt values. All statistical analyses were performed using R version 4.2.0 (R Core Team, 2022).

## Supplemental item titles and legends

**Figure S1.**
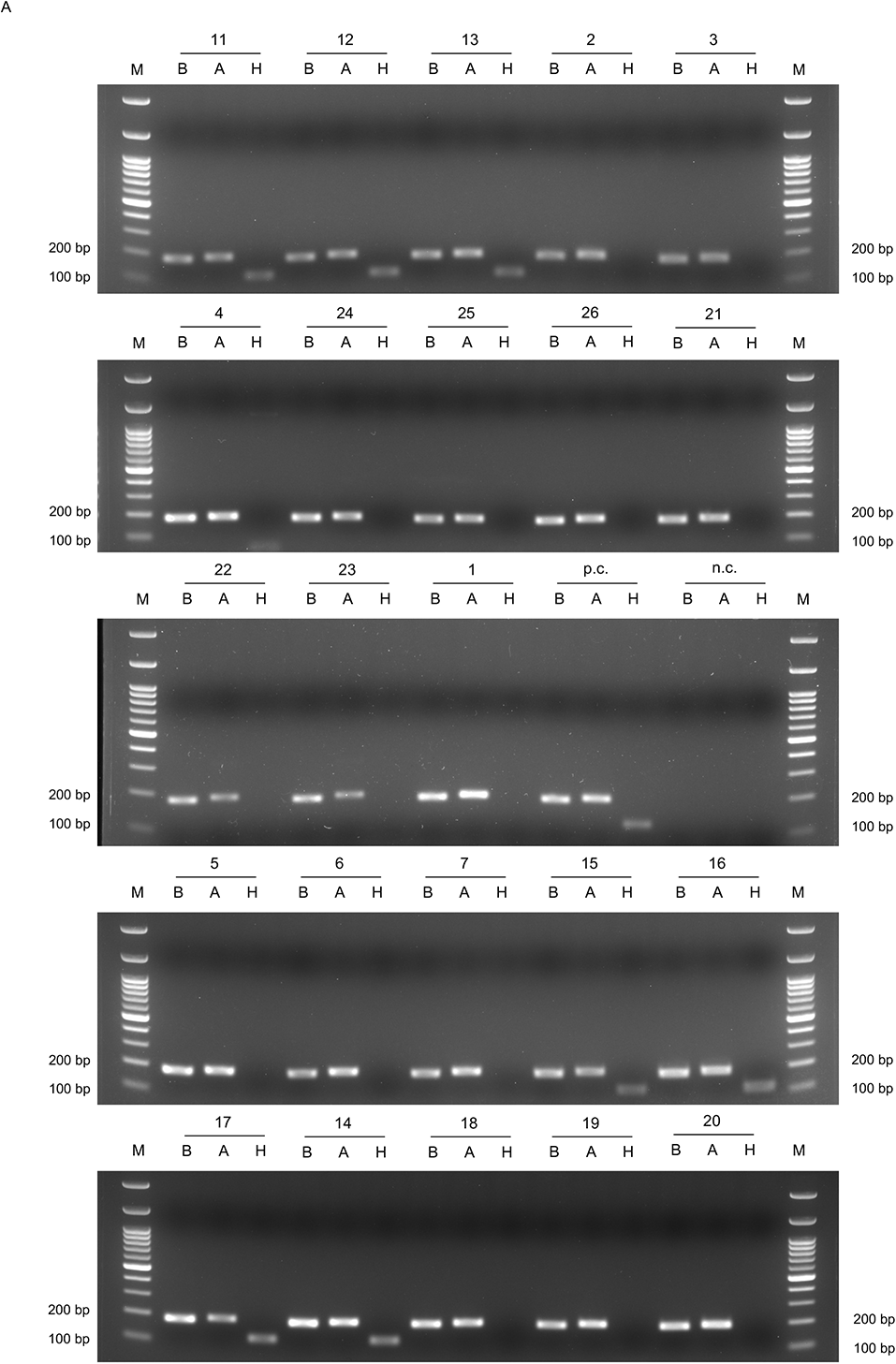
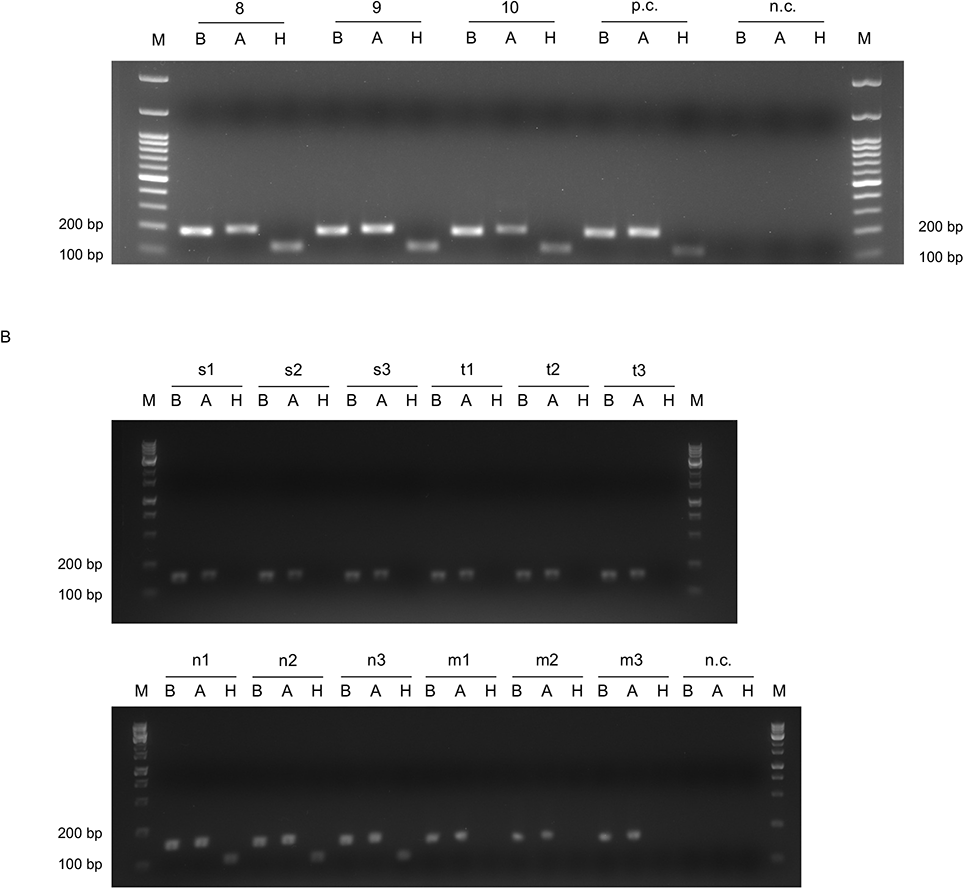
Diagnostic PCR detection of *Buchnera*, *Arsenophonus*, and *Hamiltonella* of *Ce. japonica*, related to Figure 1. (A) Symbiont detection in *Ce. japonica* populations. Each sample ID number corresponds to that of Figure 1B–D. (B) Symbiont detection in *Ce. japonica* individuals. Samples of s1–3, t1–3, n1–3, and m1–3 were derived from Sakata-koen (populations #2–4 in Figure 1), Tsushima (#24–26), Norikura (#14), and Matsudaira (#18–20), respectively. B, A, H, M, p.c., and n.c. means *Buchnera*, *Arsenophonus*, *Hamiltonella*, DNA size marker, positive control and negative control, respectively. Each bacterial symbiont was detected using specific primers targeting a single-copy gene, *dnaK*. PCR product sizes of *DnaK* of *Buchnera*, *Arsenophonus*, and *Hamiltonella* are 186, 192, and 117 bp, respectively. DNA size markers shows 3,000, 1,500, 1,000, 900, 800, 700, 600, 500, 400, 300, 200, and 100 bp, from top to bottom.

**Figure S2.**
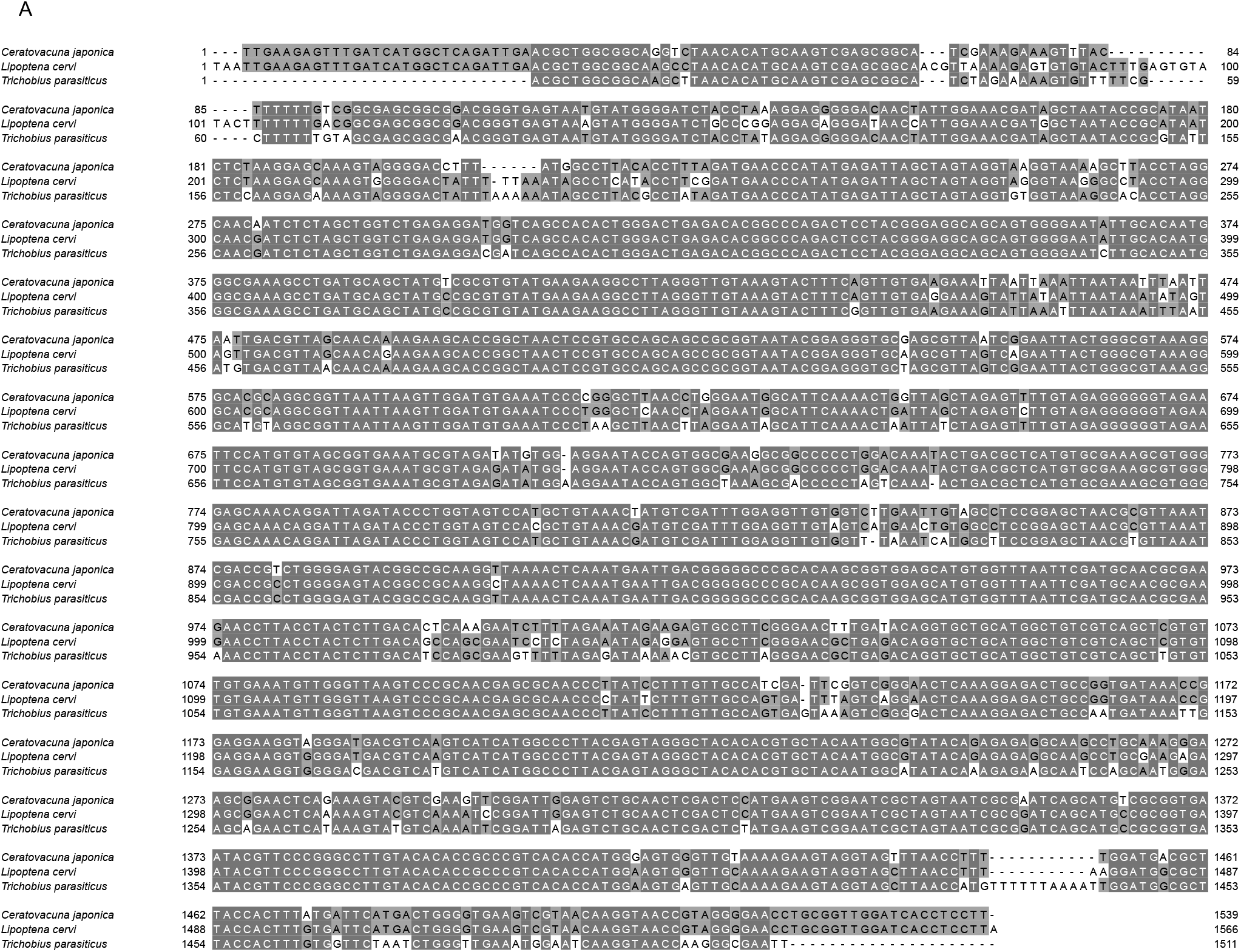
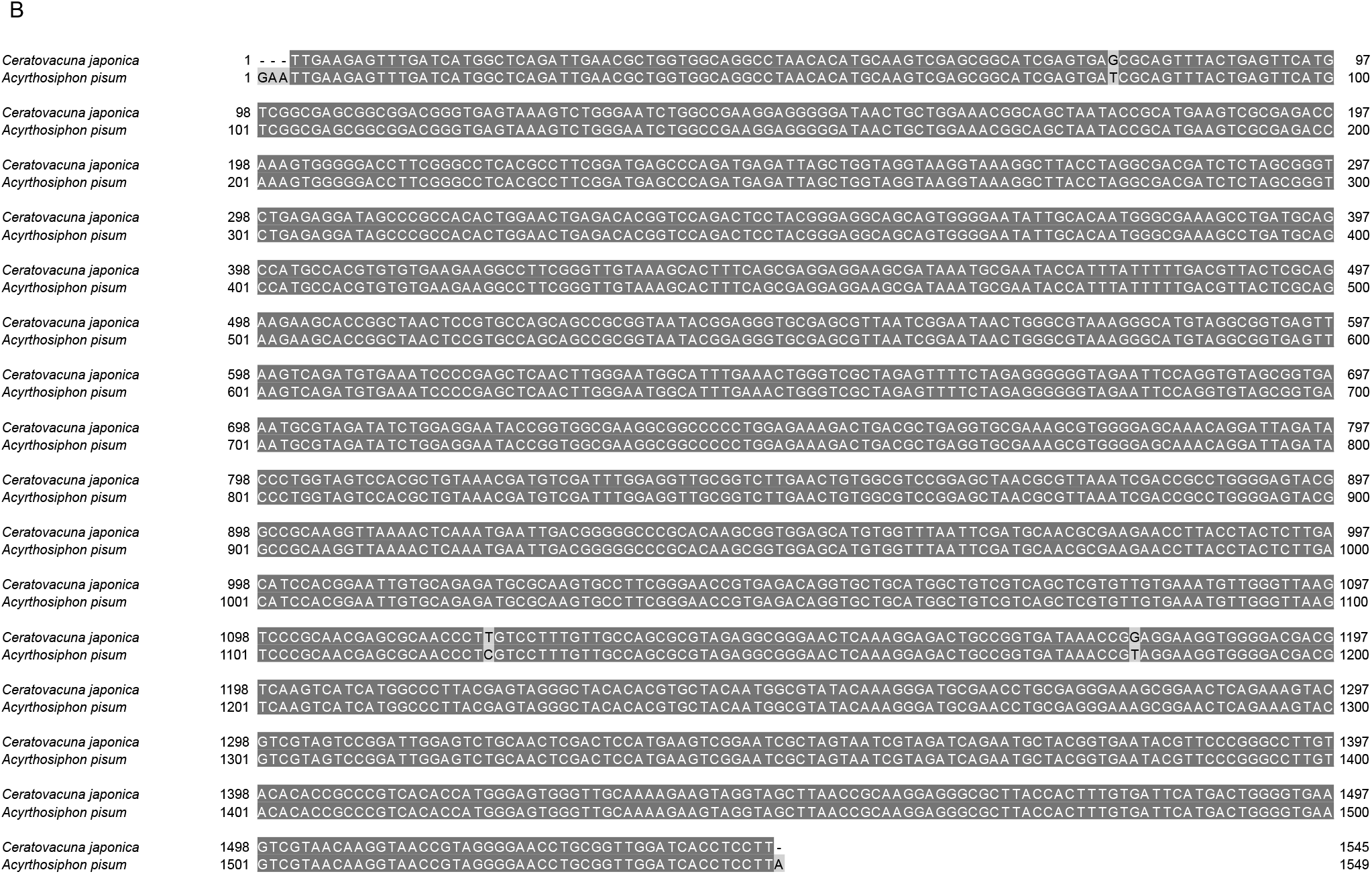
Alignments of 16S rRNA sequences of *Arsenophonus* and *Hamiltonella*, related to Figure 2. (A) A multiple alignment of *Arsenophonus* symbionts of *Ce. japonica*, *L. cervi*, and *T. parasiticus* is shown. Sequence identities of *Arsenophonus* symbionts between *Ce. japonica* and *L. cervi*, and those between *Ce. japonica* and *T. parasiticus* are 93.0% and 89.4%, respectively. (B) A pair-wise alignment of *Hamiltonella* symbionts of *Ce. japonica* and *A. pisum*. The sequence identity between these two sequences is 99.8%. See Table S3 for the accession number of 16S rRNA sequences used for these alignments.

**Figure S3.**
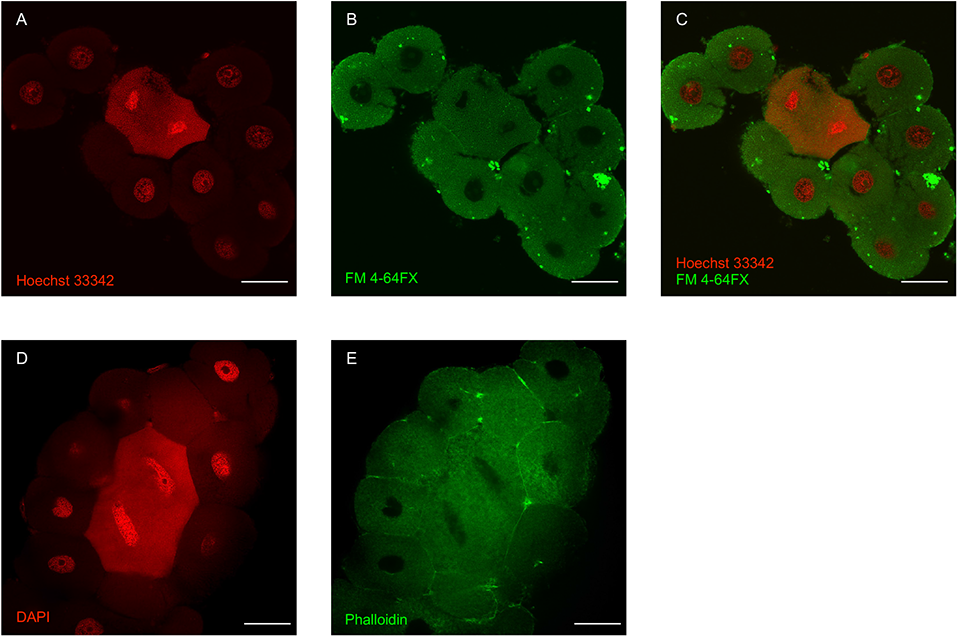
Structure of bacteriome, related to Figure 3. Confocal microscopy images of bacteriomes are shown. (A) Hoechst 33342 signals. (B) FM 4-64FX signals. (C) Merged image of (A) and (B). (D) DAPI signals. (E) Phalloidin signals. Scale bars show 50 µm in (A–E).

**Figure S4.**
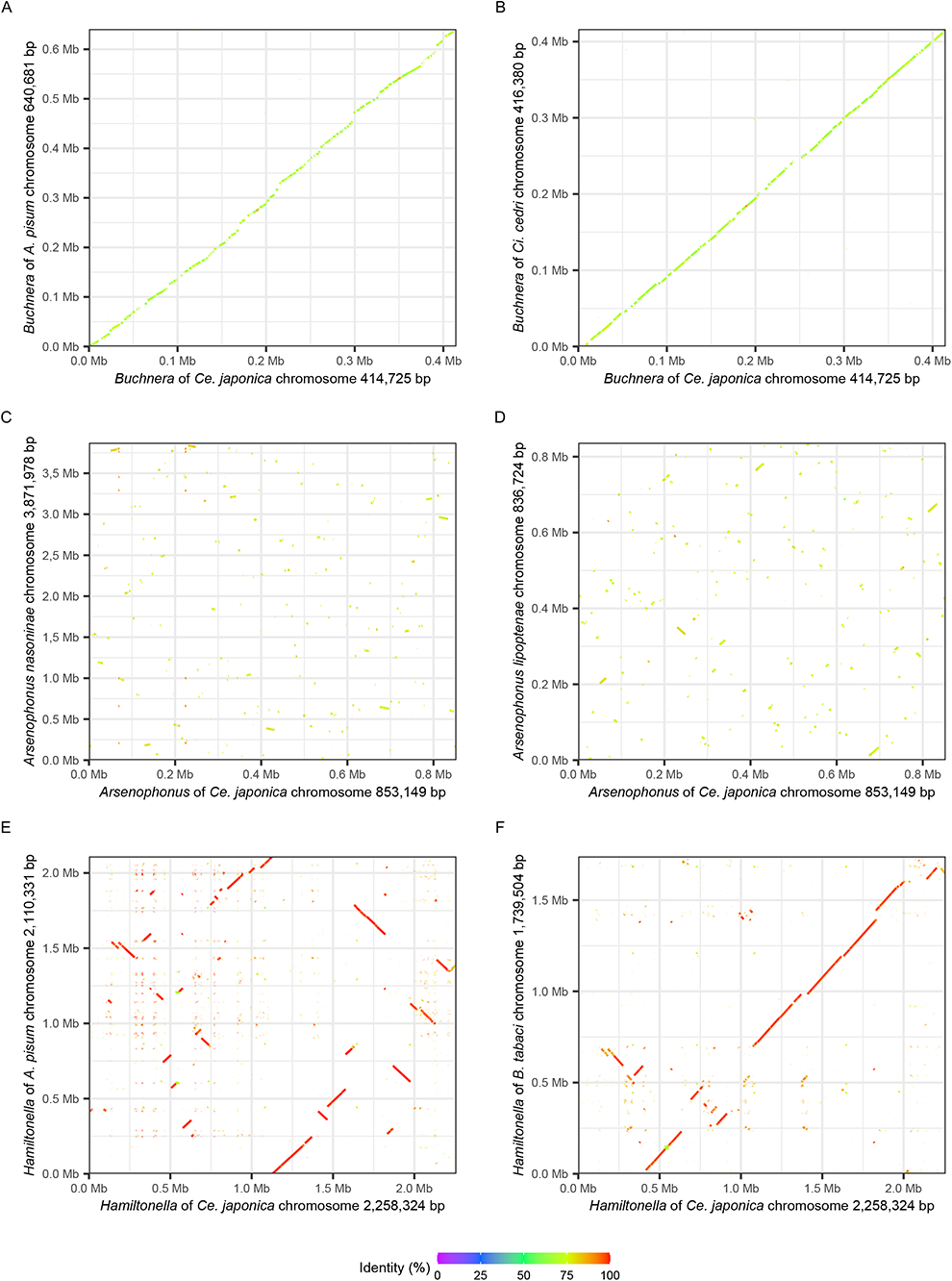
Synteny plots of chromosomes of *Buchnera*, *Arsenophonus*, and *Hamiltonella*, related to Figure 4. Genomic synteny between symbionts is visualized. (A) *Buchnera* of *Ce. japonica* and *A. pisum* (BA000003.2). The synteny is well conserved among the two *Buchnera* genomes, whereas several gaps are discernible. (B) *Buchnera* of *Ce. japonica* and *Ci. cedri* (CP000263.1). The synteny is well conserved, whereas several gaps are discernible. (C) *Arsenophonus* of *Ce. japonica* and *N. vitripennis* (CP038613.1). Little syntenic blocks are recognizable. (D) *Arsenophonus* of *Ce. japonica* and *L. cervi* (CP013920.1). Little syntenic blocks are recognizable. (E) *Hamiltonella* of *Ce. japonica* and *A. pisum* (CP001277.1). Syntenic blocks are observed with a number of inversions and rearrangements. (F) *Hamiltonella* of *Ce. japonica* and *B. tabaci* (CP016303.1). The synteny looks well conserved with some deletions and inversions.

**Figure S5.**
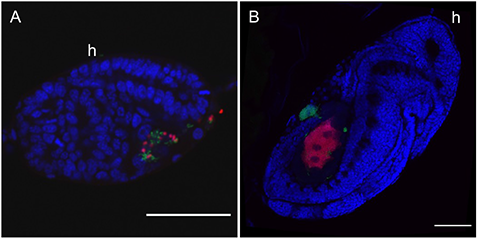
Infection process of *Hamiltonella* during the embryogenesis, related to Figure 5. (A) The developmental stage of S-shape embryo. *Hamiltonella* is also infected into the embryo from the posterior part with *Arsenophonus*. (B) The developmental stage of early germ band retraction after katatrepsis. Masses of *Hamiltonella* are observed around the bacteriome. (A and B) Blue (DAPI), green (Cy5) and red (Cy3) indicates nuclei, *Hamiltonella*, and *Arsenophonus*, respectively. Scale bars show 50 µm in (A and B).

**Table S1. The abundance of bacterial symbionts by V1–V2 region amplified from 16S rDNA in *Ce. japonica* and the sample information, related to** Figure 1.

**Table S2. The abundance of bacterial symbionts by V3–V4 region amplified from 16S rDNA in *Ce. japonica* and the sample information, related to** Figure 1.

**Table S3. 16S rRNA sequence information used for phylogenetic analyses and alignments, related to** Figure 2 **and S2.**

**Table S4. Predicted genes and annotations of *Buchnera* CJ genome, related to** Figure 4.

**Table S5. Predicted genes and annotations of *Arsenophonus* CJ genome, related to** Figure 4.

**Table S6. Predicted genes and annotations of *Hamiltonella* CJ genome, related to** Figure 4.

**Table S7. Lipid A biosynthesis genes in symbiotic bacteria, related to** Figure 4.

Cj: *Ceratovacuna japonica*, Ap: *Acyrthosiphon pisum*, Lc: *Lipoptena cervi*, Ac: *Aphis craccivora*, Nv: *Nasonia vitripennis*, Ts: *Tuberolachnus salginus*, Cc: *Cinara cedri*. *Buchnera* of *Ac. pisum* (GCA_000009605.1); *Arsenophonus* of *L. cervi* (GCA_001534665.1); *Arsenophonus* of *Ap. craccivora* (GCA_013460135.1); *Arsenophonus* of *N. vitripennis* (GCA_004768525.1); *Serratia* of *T. salginus* (GCA_900016775.1); *Serratia* of *Ci. cedri* (GCA_000238975.1); *Serratia* of *Ac. pisum* (GCA_008370165.1); *Serratia marcescens* (GCA_000336425.1); *Blochmannia floridanus* (GCA_000043285.1); *Wigglesworthia glossinidia* (GCA_000008885.1); *Baumannia cicadellinicola* (GCA_000013185.1); *Escherichia coli* (GCA_000005845.2). Φ: pseudogenes

